# A metabolic atlas of the *Klebsiella pneumoniae* species complex reveals lineage-specific metabolism that supports co-existence of diverse lineages

**DOI:** 10.1101/2024.07.24.605038

**Authors:** Ben Vezina, Helena B. Cooper, Martin Rethoret-Pasty, Sylvain Brisse, Jonathan M. Monk, Kathryn E. Holt, Kelly L. Wyres

## Abstract

The *Klebsiella pneumoniae* species complex inhabits a wide variety of hosts and environments, and is a major cause of antimicrobial resistant infections. Genomics has revealed the population comprises multiple species/subspecies and hundreds of distinct co-circulating sub-lineages that are associated with distinct gene complements. A substantial fraction of the pan-genome is predicted to be involved in metabolic functions and hence these data are consistent with metabolic differentiation as a driver of population structure. However, this has so far remained unsubstantiated because in the past it was not possible to explore metabolic variation at scale.

Here we used a combination of comparative genomics and high-throughput genome-scale metabolic modelling to systematically explore metabolic diversity across the *K. pneumoniae* species complex (n=7,835 genomes). We simulated growth outcomes for each isolate using carbon, nitrogen, phosphorus and sulfur sources under aerobic and anaerobic conditions (n=1,278 conditions per isolate). We showed that the distributions of metabolic genes and growth capabilities are structured in the population, and confirmed that sub-lineages exhibit unique metabolic profiles. *In vitro* co-culture experiments demonstrated reciprocal commensalistic cross-feeding between sub-lineages, effectively extending the range of conditions supporting individual growth. We propose that these substrate specialisations promote the existence and persistence of co-circulating sub-lineages by reducing nutrient competition and facilitating commensal interactions via negative frequency-dependent selection.

Our findings have implications for understanding the eco-evolutionary dynamics of *K. pneumoniae* and for the design of novel strategies to prevent opportunistic infections caused by this World Health Organization priority antimicrobial resistant pathogen.

## Introduction

*Klebsiella pneumoniae* is a ubiquitous Gram-negative bacterium and major cause of opportunistic healthcare associated infections with significant global burden (1). Novel prevention and control strategies are urgently required to target this organism, in particular the growing numbers of multi-drug resistant strains that cause infections that are extremely difficult to treat (2). However, our capacity to design effective control measures is complicated by the extraordinary diversity among *K. pneumoniae* and gaps in our knowledge about how key types of variation such as metabolic variation, intersect with ecology and population structure.

Isolates identified as *K. pneumoniae* in clinical laboratories include *K. pneumoniae sensu stricto* in addition to six other taxa (species and sub-species) collectively known as the *K. pneumoniae* species complex (*Kp*SC) (3). These species are separated by 3-4% nucleotide divergence in their core genes, and each is further subdivided into hundreds or thousands of phylogenetically distinct sub-lineages separated by ∼0.5% nucleotide divergence (3, 4), but with access to a highly diverse shared gene pool. It is well known that gene content differences play a role in driving differential sub-lineage epidemiologies i.e. where a subset are recognised as globally-distributed multi-drug resistant agents of healthcare-associated infections or as ‘hypervirulent’ community-acquired pathogens (usually drug-susceptible) (3). It has also been shown that total gene content is non-randomly distributed between sub-lineages and that up to 37% of the total *Kp*SC gene pool encodes proteins contributing to cellular metabolism (4).

We therefore hypothesised that sub-lineages differ in terms of metabolic capabilities, which likely contributes to their epidemiological behaviours and facilitates the maintenance of diverse sub-lineages within the population. A similar hypothesis was proposed for the Gram-positive bacterial pathogen, *Streptococcus pneumoniae* (5), and is consistent with genomic studies of other organisms that have implicated a non-random distribution of metabolic traits within the population (6–10). However, the complexities of cellular metabolism make systematic prediction of phenotypes from genome data difficult, limiting capacity for large-scale analyses of population metabolism.

Genome-scale metabolic models attempt to encapsulate the total metabolic potential of an organism and can be used to predict metabolic phenotypes (11). Comparative metabolic modelling analyses can be utilised to probe diversity within bacterial species populations, and recent larger-scale studies (n = 50-2,446 isolates) have highlighted the power of these approaches to identify industrially relevant, ecology-, drug-resistance- and pathogenicity-associated metabolic traits (12–17). While a recent study of 2,773 metabolic models confirmed that *E. coli* phylogroups (which are approximately as diverse as *Kp*SC species) are associated with unique metabolic reactions (18). However, none of these works have attempted to investigate the interplay between phenotypic diversity and fine-scale population structure i.e. to understand how diversity is distributed within and between phylogenetic sub-lineages. Such information is crucial to understanding the interaction between metabolism, ecology and evolution within species, and for design of broad-acting control strategies that seek to exploit metabolic vulnerabilities.

Here, we describe a population-level analysis of 7,835 *Kp*SC genomes to evaluate the distribution and diversity of metabolism in the context of the population structure, first at the level of species, then sub-lineages within species. We performed quantitative analyses of metabolic gene content, and constructed metabolic models from which we predicted 1,278 metabolic phenotypes for each isolate. This approach allowed us to directly infer the phenotypic impact of genetic variation and explore phenotype variation within a single species in the context of the established *Kp*SC population genomics framework (3) and at a scale that was not previously possible. We validated a subset of these phenotypes via experimental assays. Our data reveal considerable variation between and within species that is non-randomly distributed across sub-lineages. Notably, we show for the first time that *Kp*SC sub-lineages, including multi-drug resistant and hypervirulent clones, are associated with distinct substrate usage profiles that facilitate commensal interactions and likely promote co-existence.

## Materials and Methods

### Genome acquisition, quality filtering and genotyping

Genomes were included from 29 medium-large-scale studies (4, 19–45) that comprised diverse KpSC collections (**Table S1**). Outbreak investigations were explicitly excluded to reduce clonal oversampling and genomes were de-replicated to minimise sampling biases for overrepresented sub-lineages (see **Supplemental Methods**). Reads were acquired from the European Nucleotide Archive using the enaDataGet and enaGroupGet scripts from enaBrowserTools v1.6 (46), trimmed with Trim Galore v0.5.0 using -q 20 option (47) then assembled with Unicycler v0.4.7 with the --keep 0 option (48). Assemblies failing the established KlebNET-Genomic Surveillance Platform inclusion criteria were excluded (>500 contigs, total length <4,969,898 or >6,132,846 bp). Graphical Fragment Assembly (GFA) files were analysed to count GFA dead ends using getUnicyclerGraphStats.py from Unicycler (48)

Assemblies were annotated with Bakta v1.1.1 (49) with the ‘--gram –‘ option and Bakta database (date accessed: 1/09/2021), while Kleborate v2.0.4 (50) was used to identify species and Sequence Types (STs). Pathogenwatch (https://pathogen.watch/) and BIGSdb (https://bigsdb.pasteur.fr/) (51, 52) were used to assign LIN codes (53) and sub-lineages. Pairwise ANI was calculated using FastANI v1.33 (54) using --fragLen 3000.

### Pangenome analysis

The 7,835 filtered and dereplicated *Kp*SC genomes were split into species, then Panaroo v1.2.8 (55) was run with the following options: ‘--clean-mode sensitive -a core --aligner mafft --no_clean_edges --core_threshold 0.98 --merge_paralogs --remove-invalid-genes’. Due to computational limits, the 6,676 *K. pneumoniae* genomes were split into two groups of 3,338 (based on closest genetic distance) and run separately on Panaroo. The output graphs for all species were then merged using the panaroo-merge command with the following options: ‘--merge_paralogs’ to obtain the final pangenome.

### Metabolic gene identification and analysis

To analyse metabolic pathways, the pan_genome_reference.fa file from Panaroo was translated using the transeq tool from EMBOSS v6.6.0 (56) with the following options: ‘-table 11 -frame 1’. Translated sequences were analysed using kofam_scan v1.3.0 (https://github.com/takaram/kofam_scan), the command line version of KEGG’s (57) kofamKOALA (58). KEGG release 94.0 ver. 2020-04-02.

Genes with an e-value of ≤0.001 were retained and analysed using KEGG Mapper (https://www.genome.jp/kegg/mapper/reconstruct.html) to identify pathway information. Using the BRITE tab, genes which fell under the ‘metabolism’ subheading were used to classify genes as metabolic or non-metabolic. KEGG ortholog assignments were used to convert the Panaroo gene presence/absence matrix into a metabolic ortholog matrix. This allowed grouping of functional isozymes which would otherwise have an identical KEGG function. Pairwise metabolic ortholog Jaccard similarities were calculated using the vegdist function from R package vegan v2.5-7 (59).

Twilight (60) was used to classify distribution of metabolic orthologs among *Kp*SC taxa and 48 common *K. pneumoniae* sub-lineages (n≥15 genomes each, defined by LIN codes) using the -s 5 and -s 20 commands, respectively. ‘Core’: ≥95% prevalence. ‘Intermediate’: <95% and >15% prevalence. ‘Rare’: ≤15% and >0% prevalence. ‘Absent’: 0% prevalence.

### Metabolic modelling

Draft metabolic models were generated via Bactabolize v1.0.1 (61) with the *Kp*SC-pan v2.0 model (62) as the input reference (see **Supplemental Methods**). Flux balance analysis as implemented in Bactabolize v1.0.1 was used to predict binary growth phenotypes for all possible carbon, nitrogen, phosphorus and sulfur sources supported by the reference model as sole sources in M9 minimal media, aerobic and anaerobic conditions (**Supplemental Methods**). Pairwise Jaccard similarities and population distribution of positive growth phenotypes were calculated as described above for metabolic orthologs.

To evaluate co-occurrence of metabolic traits and control for population structure, genomes were randomly subsampled to a maximum of 10 per sub-lineage (n=2,427 genomes), then predicted aerobic growth phenotypes analysed using Coinfinder v1.2.0 (63) with the -- associate option.

### *In vitro* growth experiments

To validate substrate usage predictions from the metabolic models, 13 isolates (**Table S2**) were selected from our in-house collections (4, 25, 64) and tested in triplicate on each of nine substrates in minimal media, aerobic conditions. Two experiments were performed: i) endpoint OD_600_ at 24- and 48-hours; ii) 3-day substrate coaxing to induce gene expression (65) for predicted false-negative growth results in the endpoint experiment. Isolates were coaxed onto the testing substrate via subculturing into progressively lower amounts of D-glycerol, which all isolates can use as a carbon source (10 mM on subculture 1, 5 mM on subculture 2, 0 mM on subculture 3). See **Supplemental Methods** for details.

### Co-culture experiments

We selected three pairs of sub-lineages and three corresponding pairs of substrates (four total substrates) for which one was predicted as a sub-lineage specific core trait in sub-lineage A and the other predicted as a sub-lineage specific core trait in sub-lineage B. For each sub-lineage, three isolates were selected from our in-house collections (4, 25, 66) and treated as biological replicates (**Table S3**).

Isolate pairs were grown in single and co-culture using 0.4 µm pore Transwells (Sigma) and 24-well plates (Costar, Corning). See **Supplemental Methods** for further details. Co-culture results were compared to single cultures and log_2_-fold change OD_600_ values calculated.

### Co-culture simulations

*In silico* co-culture experiments were performed using MICOM v0.32.2 (67) with the relevant *K. pneumoniae* metabolic models. Community models were built using the build function, then communities were grown in growth media matching co-culture experiments (M9 minimal media with various carbon sources) with tradeoff set to 0.5. Metabolite and reaction flux was then analysed to determine metabolite sharing from prototrophs to auxotrophs.

### Data visualisation and code availability

Statistical analysis and graphical visualisation were performed using R v4.0.3 (68), RStudio v1.3.1093 (69), with the following software packages: tidyverse v1.3.1 (70), viridis v0.5.1 (71), RColorBrewer v1.1-2 (72), ggpubr v0.4.0 (73) ggpmisc v0.4.4 (74), aplot v0.1.6 (75), colorspace v2.0-2 (76), ggtree v2.4.1 (77), vegan v2.5-7 (59), rstatix v0.7.0 (78) and ggnewscale v0.4.5 (79).

All code used to generate figures, simulations and perform statistical analysis can be found at (https://dx.doi.org/10.6084/m9.figshare.24503737).

## Results

### Metabolic traits are bimodally distributed across the population

The study dataset included 7,835 high-quality *Kp*SC genomes covering all seven taxa in the complex (**Fig. 1A**) and 1,931 sequence types (STs) (**Table S1**). Pan-genome analysis (55) identified 61,595 gene clusters, 11,184 (18.2%) of which were matched to a total of 2,601 unique KEGG Orthologs (57) with associated substrate-reaction data. This represented a lower bound estimate of the proportion of ‘metabolic genes’ in the *Kp*SC pan-genome, as 38.9% of gene clusters could not be assigned functional annotations (hypothetical proteins). Metabolism-associated KEGG Orthologs (hereafter ‘metabolic orthologs’) were bimodally distributed (**Fig. 1B, Table S4**), with 1,499 orthologs (57.4%) present in ≥95% of genomes and 899 rare orthologs present in <15% of genomes. This bimodal distribution is consistent with the general distribution of accessory genes in the *Kp*SC (4), and other bacterial species with similarly large effective population sizes (60, 80).

**Fig. 1:**
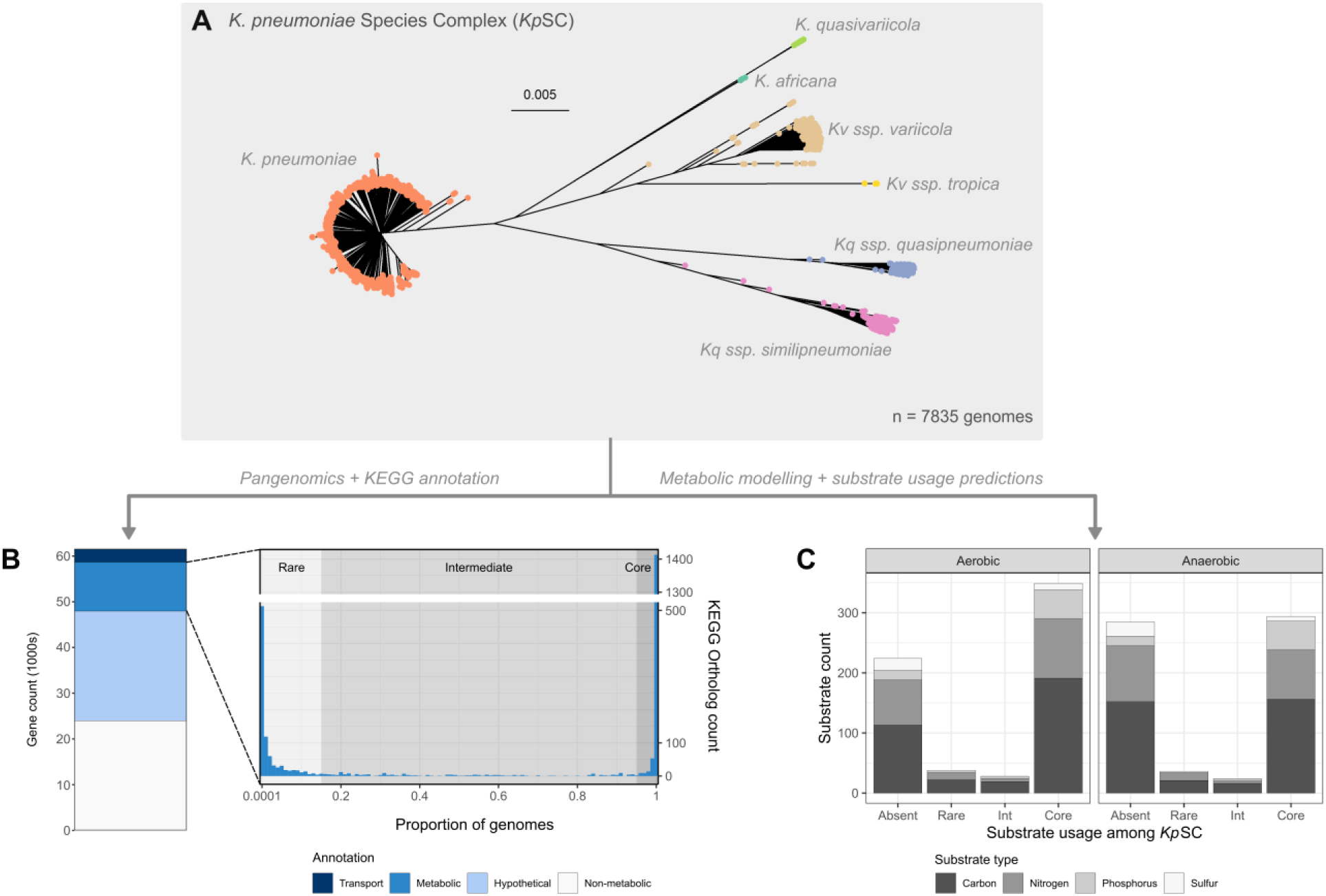
Population level metabolic diversity in the *K. pneumoniae* Species Complex. **A:** Neighbour-joining tree of the 7,835 *Kp*SC genomes used in this study. Tip colours indicate species as labelled. An interactive version can be found at https://microreact.org/project/pcTMwQZAGCsqqhbEZzaj11-a-metabolic-atlas-of-the-klebsiella-pneumoniae-species-complex. **B:** Stacked bars show pangenome content by metabolic gene (KEGG) category as indicated in legend (left). Population distribution of KEGG orthologs associated with metabolic genes (‘metabolic orthologs’, right). The light grey background shows rare orthologs (>0 to 15%), the medium grey shows intermediate orthologs (>15% to <95%) while the dark grey shows core orthologs (≥95%). **C:** Stacked bars show population frequency categories for predicted substrate usage, coloured by substrate type as indicated in legend.

To analyse the phenotypic diversity across the *Kp*SC, a strain-specific metabolic model was generated for each genome using Bactabolize (61), which was previously shown to produce models with 93.2% and 78.1% predictive accuracy for aerobic and anaerobic growth phenotypes, respectively (62). Growth phenotypes were predicted for 345 carbon, 192 nitrogen, 68 phosphorus and 34 sulfur substrates as sole sources in minimal media under aerobic and anaerobic conditions (total 1,278 conditions). Individual strain-specific models comprised a mean of 1677.0 genes ± 17.3 (standard deviation, SD) and 3330.1 reactions ± 13.5 (SD). Individual isolates were predicted to grow in a mean of 678.2 ± 17.9 (SD) conditions (**Table S5**, **Fig. S1A, Supplemental Results**).

The distribution of positive growth predictions mirrored that for metabolic gene content. At the population level, 643 conditions (47.6%) were predicted to support growth for ≥95% isolates (‘core’, **Fig. 1C**, **Table S5**) and were comprised of 349 aerobic and 294 anaerobic conditions, which systematically confirmed the facultative anaerobe status of the *Kp*SC. These 643 core conditions consisted of 207 total substrates (note that some substrates can be tested as sole sources of multiple elements e.g. carbon and nitrogen), with at least 204 of these being known human metabolites, 134 diet-related compounds and 203 gut microbiome metabolites (**Table S5**). In contrast, 509 modelled growth conditions (43.6%) did not support growth of any isolates (224 aerobic and 285 anaerobic conditions). Carbon substrate usage displayed the greatest variability, with 41 aerobic and 37 anerobic conditions predicted to support growth in <95% isolates (**Fig. 1C, Fig. S1B**).

### Metabolic variation is structured within and between species

The set of core growth conditions for each *Kp*SC taxon was greater than the species complex as a whole (11-39 additional conditions each). Each taxon was associated with a distinct core growth profile mirrored by distinct core metabolic ortholog profiles as expected (66, 81) (**Fig. S2 and S3, Tables S4 and S5, Supplemental Results**), and there was evidence of specialisation at the level of macromolecular functions as determined by COG analysis (**Fig. S4, Table S1, Supplemental Results**). These predicted growth capabilities were generally consistent with biochemical tests described in formal species definitions (82–85). However, there were a small number of notable discrepancies that indicate that species identification protocols should be revisited (see **Supplemental Results** and **Table S6** for details).

While individual *Kp*SC taxa are clearly defined by distinct metabolic profiles, there was also clear evidence of intra-taxon variation (**Fig. S2 and S3**). Four taxa were represented by sufficient genomes for comparisons (n>200 each); *K. pneumoniae* (hereafter *Kp,* n=6,652), *Klebsiella quasipneumoniae* subsp. *quasipneumoniae* (*Kqq*, n=201), *K. quasipneumoniae* subsp. *similipneumoniae* (*Kqs*, n=285) and *Klebsiella variicola* subsp. *variicola* (*Kvv,* n=672). In *Kp,* 938 metabolic orthologs were variably present in <95% genomes (544, 586 and 738 orthologs variable among *Kqq, Kqs* and *Kvv*, respectively) and 79 conditions were predicted to support variable growth (81, 78 and 67 for *Kqq, Kqs* and *Kvv*). We hypothesised that this variation is not randomly distributed in the population, and our analysis of all-vs-all genome pairs within taxa showed statistically significant positive relationships between pairwise Average Nucleotide Identity (ANI, generally indicative of ancestral relatedness) and pairwise Jaccard similarity across metabolic genes (general linear hypothesis test, p<2×10^−16^ for each of the four well sampled taxa, **Fig. 2A**). Comparison of the regression gradients indicated significant differences between taxa; however, the R^2^ values were low (range 0.04-0.48), indicating a large amount of unexplained variation.

**Fig. 2:**
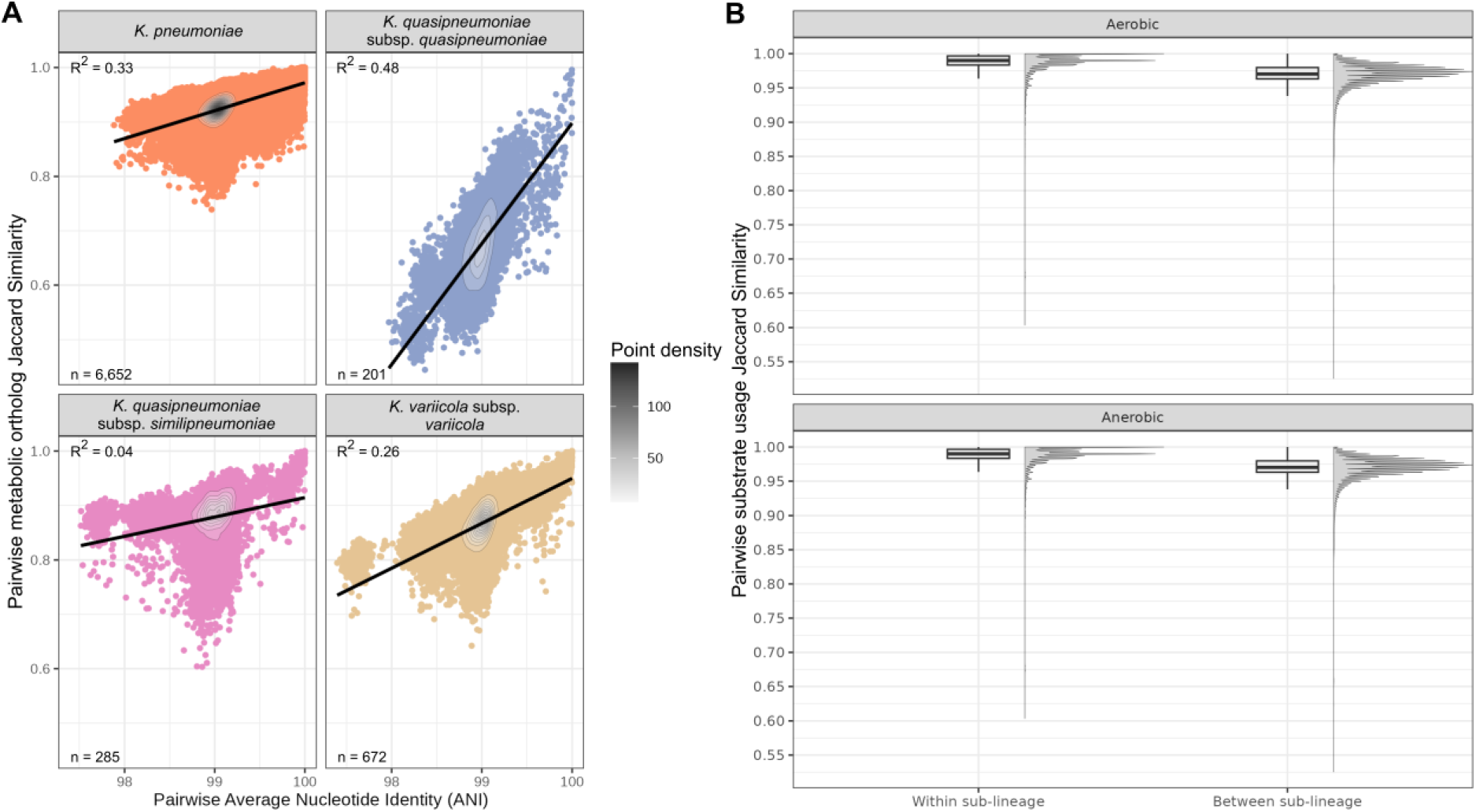
Metabolic heterogeneity within taxa is non-randomly distributed. **A:** Pairwise Average Nucleotide Identity (ANI) vs pairwise Jaccard similarity of metabolic orthologs within each of the well sampled taxa (n>200 genomes). **B:** Distributions of pairwise (all vs all) Jaccard Similarity of simulated substrate usage for pairs of genomes within vs between sub-lineages in the same species. Significance asterisks excluded as each group was significantly different from every other group (p < 2×10^−16^), calculated using a non-parametric Kruskal–Wallis test with Holm correction, followed by Dunn’s *post-hoc* test.

Next we used the recently established core-genome Life Identifier Number (LIN) code scheme to assign genomes to sub-lineages (SL) (53). These sub-lineages represent deep-branching phylogenetic clades, separated by hundreds of years of evolutionary divergence (with notable exceptions discussed below). Of the 7,835 genomes, 7,833 were assigned to 1,418 distinct sub-lineages and two were unassigned due to insufficient alleles detected.

Pairwise Jaccard similarities for metabolic ortholog profiles were significantly higher within sub-lineages (median 0.956, IQR 0.947-0.965) than between sub-lineages within the same species (median 0.923, IQR 0.914-0.932), p < 0.0001 (Kruskal–Wallis test). Mirroring this, pairwise Jaccard similarities for predicted substrate usage profiles were significantly higher for pairs within sub-lineages (median 0.990, IQR 0.983-0.997) than between sub-lineages (median 0.97, IQR 0.96-0.98) for both aerobic and anaerobic substrate usage (p < 0.0001 for all comparisons, Kruskal–Wallis test with Holm correction, followed by Dunn’s *post-hoc*, **Fig. 2B**).

Finally, we sought to explore structuring among accessory metabolic traits, using a co-occurrence analysis. Twelve pairs of co-occurring substrate usages were identified and 11 of these pairs were mechanistically-linked e.g. the same substrate tested as different element sources (n=5 pairs) or substrates for which the catabolic pathways are overlapping (n=4 pairs, see **Supplemental Results**). Just one co-occurring phenotype pair (formamide and Fe(III)dicitrate as sources of carbon) was not obviously mechanistically linked, indicating that they may be co-selected or encoded by genes that are physically linked in the genome or on plasmids. Accordingly, we have identified the genes enabling use of formamide and Fe(III)dicitrate as sources of carbon co-located on plasmids in the completed genomes of multiple unrelated strains e.g. INF008 (SL221), INF044 (SL258) and INF119 (SL17).

### Major sub-lineages are defined by unique metabolic profiles

To explore the association between metabolism and population structure at higher-resolution, we calculated the relative prevalence of predicted growth phenotypes within 48 *Kp* sub-lineages that were each represented by genomes from broad geographies and time-spans (≥15 genomes, 3-12 geographic regions, 5-86 years). These included sub-lineages corresponding to globally distributed multi-drug resistant (SL14, SL15, SL17, SL29, SL37, SL101, SL147, SL258, SL307) and hypervirulent clones (SL23, SL25, SL65, SL86) (3). The data showed that each sub-lineage is associated with a distinct, core metabolic fingerprint, indicating metabolic specialisation (**Fig. 3**, **Fig. S5**, **Table S5**), with 253-316 core aerobic and 198-223 core anaerobic conditions each. Note that in some cases the sub-lineage specific core was smaller than the *Kp* species core reported above using the ≥95% genome threshold (363 aerobic, 307 anaerobic conditions), because most sub-lineages represent far less than 5% of the total *Kp* genome collection. This contradiction highlights the influence of sample size and diversity on core definitions, which should be considered as indicative of the general trends in the population. No pair of sub-lineages shared the same core growth profile, even pairs that are thought to share relatively recent common ancestry such as SL14 and SL15 (common ancestor approximately 45 years ago (4)) and SL65 and SL25 (common ancestor yet to be dated) (86) (**Fig. 3**, **Fig. S5**). However, there was also clear evidence of diversity within sub-lineages, with 3-102 variable growth phenotypes each, a pattern that was replicated by metabolic ortholog distributions (**Fig. S3, Fig. S5, Supplemental Results**).

**Fig. 3:**
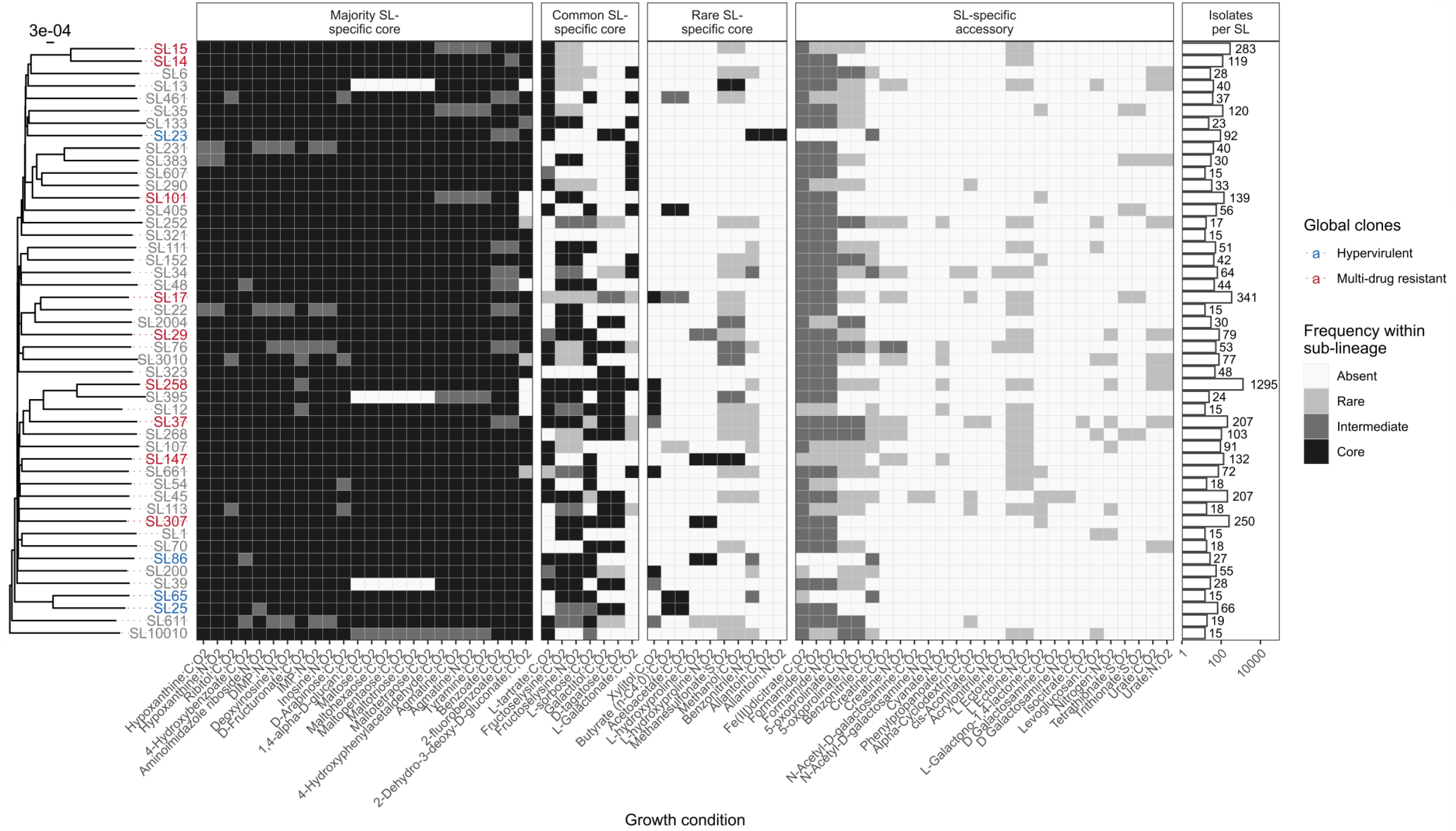
Distinct metabolic fingerprints of sub-lineages within species. Heatmap showing frequency of variable aerobic substrate usage across 48 global *K. pneumoniae* sub-lineages (SL). Rows are ordered by phylogeny. Sub-lineage labels are coloured to indicate the globally-distributed clones described in (3): Blue shows hypervirulent while red shows multidrug resistant. Substrate shown along X-axis. The element source of each substrate is semi-colon separated and abbreviated for brevity: C = Carbon, N = Nitrogen and S = Sulfur. O2 indicates aerobic conditions. Frequency of substrate usage indicated by shading as shown in legend. Number of isolates per each sub-lineage shown in bars. Anaerobic growth predictions shown in **Fig. S5.**

After excluding growth conditions predicted to support growth among ≥95% (46/48) *K. pneumoniae* sub-lineages (n=615 conditions, 340 aerobic and 275 anerobic, corresponding to 202 total substrates) and those that did not support any growth among the 48 sub-lineages (n=525 conditions), the data support a broad classification of phenotypes as follows: i) 89 that were core in some sub-lineages (range=1-45 sub-lineages) but variably present in others (range=1-17) which we call ‘sub-lineage-specific core traits’; ii) 13 that were not core in any sub-lineage (accessory in 3-46 sub-lineages, including the only pair of co-occurring phenotypes that was not mechanistically linked) (**Fig. 3**, **Fig. S5**). The sub-lineage-specific core traits could be further divided into those that were core to the majority (≥66%) of sub-lineages (n=57 phenotypes that we call ‘majority sub-lineage-specific core traits’), those that were core to an intermediate number of sub-lineages (≥25% to <66%, n=13 phenotypes, corresponding to six substrates, that we call ‘common sub-lineage-specific core traits’) and those that were core in only very few sub-lineages (<25%, n=13 ‘rare sub-lineage-specific core traits’, eight aerobic and five anerobic, corresponding to seven total substrates). We propose that the majority sub-lineage specific core traits are unlikely to play a major role in defining sub-lineage specific evolutionary trajectories or ecological interactions, and hence we focussed our follow-up investigations on the common and rare sub-lineage specific core traits.

We have previously confirmed predictive accuracies ≥97% for five of the six common sub-lineage-specific core substrates; L-sorbose, L-tartrate, galactitol, D-tagatose and L-galactonate as sole sources of carbon in aerobic conditions (62) (tested using the Omnilog phenotyping microarray). The sixth, fructoselysine, is not present in the Omnilog array and could not be tested here due to its high cost. However, we tested seven of the rare sub-lineage-specific core growth substrates under aerobic conditions, confirming high accuracy for L-hydroxyproline as a carbon source (84.6%) and variable accuracies for the remainder (**Table S2, Supplemental Results**), which was not unexpected as rare substrate usage is comparatively understudied and therefore difficult to predict.

### Sub-lineages co-operate through cross-feeding metabolites

We have shown substrate usage varies across the *Kp*SC and some of this variation is sub-lineage-dependent. We hypothesised that such diversity promotes the maintenance of distinct sub-lineages by limiting inter-lineage competition and/or promoting commensalism. We tested this hypothesis using *in vitro* co-culture experiments.

We selected four carbon substrates (galactitol, L-tartrate, L-sorbose, L-hydroxyproline) for use in three independent substrate combinations and corresponding sub-lineage pairs for which predicted substrate-usages were mutually exclusive sub-lineage-specific core traits. One isolate from each sub-lineage was selected for co-culture aerobically in each of two conditions (minimal media plus each of the sub-lineage-specific core substrates), such that each experiment contained one isolate that was able to utilise the substrate as a sole carbon source (the prototroph) and one that was not (the auxotroph). In short, Strain A as an auxotroph and Strain B as a prototroph, after which we then swapped their roles via use of another substrate, so that A would be a prototroph and B would be an auxotroph. This was done within each pair to evaluate reciprocal cross-feeding between sub-lineages under different conditions. Growth of the auxotroph was measured using OD_600_ and compared to the same isolate growing in monoculture in the same media. Co-cultured isolates were physically separated by a Transwell membrane but had shared access to the growth medium (**Fig. 4A**). A unique aspect of this experiment was the use of independent isolates from the same sub-lineages as biological replicates rather than replicates of the same isolate. This allowed us to limit the impact of strain-specific growth biases while demonstrating conserved behaviour across a greater number of independent isolates. Sub-lineages representing clinically-relevant clones were selected (multi-drug resistant SL37 vs hypervirulent SL29, hypervirulent SL86 vs hypervirulent SL23 and multi-drug resistant SL17 vs SL29).

**Fig. 4:**
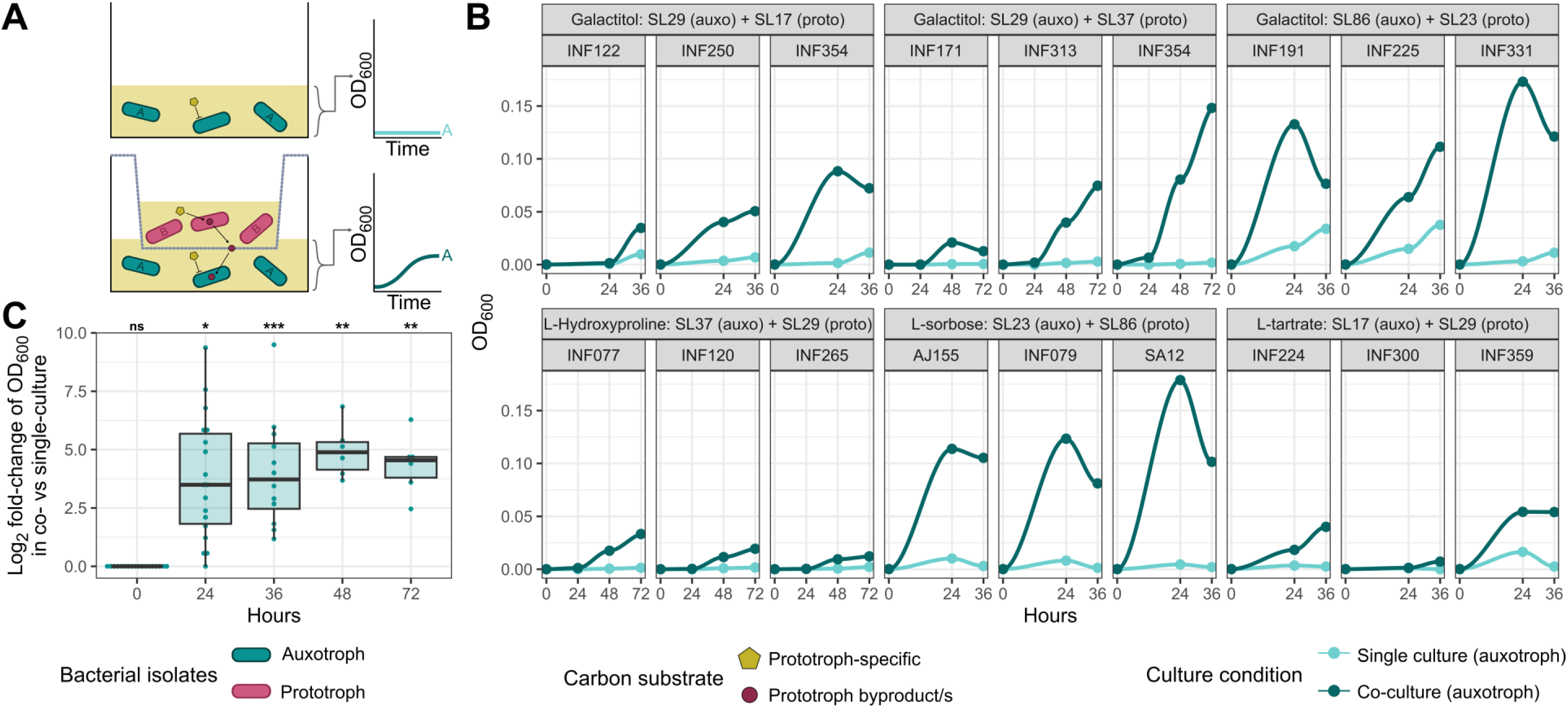
Nutrient prototrophic lineages support growth of auxotrophic lineages. Co-culture growth assays between prototrophs and auxotrophs under various substrate usage conditions. Time points captured were at 0, 24, 36, 48 and 72 hours depending upon the substrate pair (**Methods**). **A:** Graphic demonstrating the experimental design measuring commensalism between an auxotrophic strain unable to utilise a carbon source and a prototrophic strain which can. The growth of the auxotroph was compared in single vs co-culture conditions. **B:** Growth results (OD600) over time for each auxotrophic strain in the bottom well (n=3 strains per sub-lineage as biological triplicates) in single and co-culture with a prototroph for the substrate. **C:** Log2-fold change distribution of growth results (OD600) at each timepoint under different culture conditions. Statistical significance comparing the OD600 values of co-culture to single culture is shown above plot (Dunn’s *post-hoc* with Holm correction, * p < 0.05, ** p < 0.01, *** p < 0.001). Raw data can be found in **Table S3**.

Our data showed that co-culturing auxotrophs with prototrophs in the presence of a sub-lineage-specific core substrate significantly increased the OD_600_ of the auxotrophs (**Fig. 4B**) with mean log_2_-fold change 3.78 ± 2.64 SD at 24 hours, p = 0.0128; 4.02 ± 2.34 SD at 36 hours, p = 0.0001; 4.94 ± 1.14 SD at 48 hours, p = 0.0039; and 4.35 ± 1.27 at 72 hours p = 0.0039; Kruskal-Wallis + Dunn’s *post-hoc* with Holm correction (**Fig. 4C**). Corresponding *in silico* co-culture simulations indicated that prototrophs were likely cross-feeding the auxotrophs by catabolising the sole carbon substrates, then exporting metabolites that were capable of supporting auxotroph growth (see **Supplemental Results**).

To confirm this phenomenon only occurred in the presence of a cross-feeding partner and eliminate the possibility of contamination or horizontal transfer of the genes conferring the prototroph phenotype, aliquots from each co-culture well were plated onto LB agar to confirm a single, distinct colony morphology. Colonies were then inoculated into M9 minimal media plus the auxotrophic substrate as a sole carbon source, confirming the original growth behaviour of each isolate (**Table S3**). Together these results indicated that cross-feeding was occurring from the prototrophs to the auxotrophs across: i) multiple genetic backgrounds (sub-lineages); ii) to and from each strain partner within each pair under different conditions; iii) across independent isolates within sub-lineages, confirming that commensal interactions can occur between co-habiting *Kp* sub-lineages.

## Discussion

Here we present a bacterial population metabolism study of the *K. pneumoniae* species complex, in which we used quantitative metabolic modelling to predict and compare growth phenotypes for >7,000 members of a single species complex, and contextualised the results against an established population genomics framework (3). This approach allowed us to identify key metabolic differences that are indicative of metabolic fingerprints at both the species and sub-species (sub-lineage) levels. Our subsequent *in vitro* co-culture experiments using multiple independent isolates from multiple sub-lineages and multiple substrate combinations (**Fig. 4**) provide evidence that these metabolic differences can facilitate commensal interactions between sub-lineages.

Our data shows that growth capabilities are structured in the population: while there was clearly variation within and between sub-lineages, each of the 48 sub-lineages we examined was associated with a distinct set of core traits (metabolic orthologs and growth phenotypes), which likely represents vertical inheritance from the sub-lineage common ancestor (**Fig. 3**, **Fig S5**). This finding is consistent with a model of negative frequency-dependent selection as a driver for the maintenance of diverse sub-lineages (entities) in the population (87, 88). Where two sub-lineages compete for limited resources, e.g. food, the ability to consume a substrate that cannot be consumed by a competitor provides a selective advantage resulting in a relative frequency increase. This in turn results in a depletion of the food resource and shift in selective pressure to favour the competitor. Although comparatively understudied, commensal interactions such as the metabolite cross-feeding implicated by our co-culture experiments (**Fig. 4**), can drive similar frequency-dependent selective shifts (88). Together, these fluctuations prevent the exclusion of sub-lineages and drive the population to a stable equilibrium of co-existence.

Niche range no doubt further complicates the eco-evolutionary dynamics within *Kp* populations e.g. where sub-lineages are exposed to fluctuating selective pressures due to transient migration between niches, and/or where sub-lineages undergo niche adaptation. Recent analyses of contemporaneous *Kp* collections have shown that sub-lineages are not equally distributed between different hosts and environmental sources, which may indicate some level of niche-adaptation (89, 90). However, the same studies also highlighted significant diversity within niches and near-identical strains from multiple sources, which is indicative of niche migration. Unfortunately, we were unable to directly test for associations between sub-lineage, metabolism and ecological niche with our convenience sample of genomes comprising primarily human-derived isolates (6,465/7,835 genomes). Future studies will be required to probe these complex multi-way interactions and to systematically test the model of negative-frequency dependent selection, which is notoriously difficult to detect (88), requiring a long-term dataset from the same location over time, and/or laboratory protocols that are beyond the scope of the current work.

While genome-scale metabolic models are powerful approaches for exploring cellular metabolism, they are not without caveats. Firstly, model construction is reliant on curated databases that are limited by current biochemical knowledge (57), and while our pan-metabolic reference model was built from a large diverse collection of strains (62), it cannot possibly capture all metabolism in the *Kp*SC population. Hence, we are likely underestimating the metabolic diversity within the highly variable *Kp*SC. Secondly, growth prediction accuracies can vary by substrate, and are generally lower for anaerobic conditions for which the metabolic processes are less well understood (62). While many of the core substrates and common sub-lineage specific core traits have been confirmed with high accuracy in previous work, our phenotypic testing showed variable accuracy for rarer substrates. Of note, our assays showed that allantoin usage, a trait previously associated with hypervirulent infections (64) and predicted by our models as a sub-lineage specific core trait in the dominant hypervirulent SL23, may in fact be core to *Kp*SC (see **Supplemental Results**). These and other inaccurate predictions highlight gaps in our current knowledge that are noted for future investigations. Finally, metabolic models do not consider the impact of sequence variations within metabolic genes nor variations in gene regulation that will also play an important role in determining metabolic phenotypes and ecological adaptation.

Notwithstanding the caveats of *in silico* modelling, our analyses clearly challenge the assumption that *Kp* performs a defined and homogenous metabolic function, and this has important implications for how we understand and model microbial communities containing this organism. There is increasing interest in gut microbiota manipulation for prevention of pathogen colonisation and subsequent opportunistic infection (91, 92), and studies on *Kp* have thus far implicated competitive exclusion via nutrient competition as the major mechanistic driver (92, 93). Notably, the *Kp* sub-lineage-specific core traits we identified include those implicated in driving competitive interactions and colonisation resistance in the mammalian gut e.g. fructoselysine and galactitol usage (94, 95). We argue that a comprehensive understanding of the population distribution of these and other metabolic traits, will greatly benefit the design of novel control strategies targeting *Kp*, particularly those harnessing microbial competitive interactions or targeting metabolic processes.

## Author Contributions

Conceptualization: BV, SB, JMM, KEH, KLW.

Methodology: BV, SB, JMM, KEH, KLW

Validation: BV.

Formal analysis: BV, HBC, KLW.

Investigation: BV, HBC. Resources: KLW, SB, KEH.

Data Curation: BV, MR-P

Writing: BV, HBC, MR-P, SB, JMM, KEH, KLW.

Visualization: BV, KLW. Supervision: SB, KLW.

Project administration: KLW.

Funding acquisition: SB, JMM, KEH, KLW.

## Data availability

All genomic data used is publicly available - accessions and genomes used can be found in **Table S1**. All raw data can be found in **Supplemental Tables**. R and MICOM code can be found at Figshare (https://dx.doi.org/10.6084/m9.figshare.24503737). Interactive phylogeny can be found at https://microreact.org/project/pcTMwQZAGCsqqhbEZzaj11-a-metabolic-atlas-of-the-klebsiella-pneumoniae-species-complex.

## Acknowledgements

This work was supported by the MASSIVE HPC facility (www.massive.org.au).

