## Supplemental materials for "A metabolic atlas of the *Klebsiella pneumoniae* species complex reveals lineage-specific metabolism that supports co-existence of diverse lineages"

### Supplemental Methods

#### Genome dereplication

We noticed that several sub-lineages were vastly overrepresented in the dataset (e.g. due to dense sampling in specific geographies or inclusion of transmission clusters within larger collections). In order to minimise the impact of these sampling biases we dereplicated the dataset using a similar approach to that described in (1). Initially, popPUNK v2.4.0 (2) was used to finely cluster isolates. The 'create-db' function was used with the following options: '--sketch-size 1000000 --min-k 15 --max-k 29 --qc-filter prune'. Then the 'fit-model' function was used with the following options: 'dbscan --ranks 1,2,3,5 --graph-weights'. Finally, the 'fit-model' function was used again, with the options: 'refine --graph-weights --unconstrained'. Additionally, the 'poppunk_visualise' function was used, with the '--distances' and '--previous-clustering' utilising the refined model fit, to output a neighbour-joining core tree.

Isolates which shared the same popPUNK cluster, isolation source and country were sorted using newfangled_untangler.py v1.0.4 (https://github.com/bananabenana/newfangled_untangler/) then dereplicated via Assembly-Dereplicator v0.1.0 (3) with the following options: '--threshold 0.0003'.

#### Metabolic modelling

Metabolic models were generated via Bactabolize v1.0.1 (4) using the *draft_model* command with the following options: '--media_type m9 --atmosphere_type aerobic --min_coverage 25 --min_pident 80'. The *Kp*SC-pan v2.0 model (5) was used as the input reference. One-hundred and sixty of 7,835 models (2.0%) required gap-filling to enable simulation of growth on M9 minimal media with glucose (median: 1 reaction added, range: 1 – 23), using flux balance analysis to optimise the most recent version of the biomass objective function 'BIOMASS_Core_Feb2022'. (Note - we have previously shown that models gap-filled to simulate growth in minimal media plus glucose can accurately predict a range of other substrate growth phenotypes (4).)

Flux balance analysis was used to predict growth phenotypes for all possible carbon, nitrogen, phosphorus and sulfur sources supported by the reference model as sole sources of carbon, nitrogen, phosphorus or sulfur in M9 minimal media, in both aerobic and anaerobic conditions. These analyses were performed using Bactabolize's *fba* command with the following option: '--fba_spec_name m9'. All growth lower bounds and default substrate sources used in this analysis are available at https://github.com/kelwyres/Bactabolize. Models were considered capable of simulating growth when the optimised biomass value was ≥0.0001.

#### *In vitro* growth experiments

Nine growth substrates were tested: seven as sole sources of carbon (methanol, L-hydroxyproline, xylitol, butyrate, acetoacetate, D-glycerol, D-glucose), one as a sole source of nitrogen (allantoin) and one as a sole source of sulfur (methanesulfonate).

For endpoint experiments, isolates were grown overnight in 2 mL M9 media + 10 mM D-glycerol. M9 media was prepared by using 1X M9 Minimal Salts (Sigma) and adding 2 mM $MgSO_4$ and 0.1 mM $CaCl_2$ after autoclaving. 1 mL of culture was pelleted at 8000 x g for 10 minutes, supernatant discarded, then centrifugally washed using 1 mL 0.9% NaCl to remove excess carbon/nitrogen/sulfur. Isolates were then diluted 1:100 using 0.9% NaCl and 5 µL sub-cultured into 96-well plates (CLS3603, Corning) containing 200 µL M9 + substrate. Substrates were tested at the following concentrations: 400 mM methanol, 20 mM L-hydroxyproline, 20 mM xylitol, 20 mM sodium butyrate, 20 mM acetoacetate, 20 mM D-glucose, 10 mM D-glycerol 20 mM methanesulfonate, 60 mM allantoin, all at pH 7.0 and 0.2 µm filter sterilised. For allantoin as a nitrogen source, custom nitrogen-negative M9 was prepared (34 mM $NaH_2PO_4$, 64 mM $K_2HPO_4$, 1 µM $FeSO_4$, 2 mM $MgSO_4$ and 0.1 mM $CaCl_2$). For methanesulfonate as a sulfur source, sulfur-negative M9 was prepared (2 mM $MgCl_2$ instead of $MgSO_4$). Isolate negative and carbon positive controls were performed for each media. Plates were incubated at 37°C, shaking at 150 RPM and optical density read every 24 hours at $OD_{600}$ absorbance using an infinite 200Pro and i-control v2.0.10.0 (Tecan). Each experiment was performed in biological triplicate. Each well was blanked with a no isolate control, then the mean used to determine growth, with a cut-off of 0.05 (limit of detection).

Coaxing experiments were performed as above, over three days. Isolates were added to media containing the tested substrate plus D-glycerol 10 mM on subculture day 1, 5 mM on subculture day 2, 0 mM on subculture day 3. Prior to sub-culturing, cells were centrifugally washed as above and 5 µL transferred to 200 µL total media. Only subcultures on day 3 were used for analysis (0 mM D-glycerol).

##### Co-culture experiments

To rule out strain-strain antagonism prior to co-culturing, a primary inoculum was streaked onto LB agar plates and grown for eight hours at 37°C. Then, primary inoculums of additional isolates were streaked perpendicularly. Plates were incubated for 16 hours at 37°C and no growth inhibition of the perpendicular streaks was observed.

Co-culture results were compared to single cultures using $OD_{600}$ of pairs of isolates (**Table S3**) that were physically separated by 0.4 µm pores using 24-well Transwells (Sigma). All cultures were grown at 37°C, 100 RPM. Substrates were used in 1X M9 media at the following concentrations: 20 mM for galactitol, L-sorbose, L-tartrate, and 5 mM for L-hydroxyproline.

M9 media plus the tested substrate was added to the Transwell (100 µL) and the main well (800 µL). Isolates were grown overnight in 1X M9 containing 10 mM D-glycerol, then centrifugally washed as above. Cultures were diluted 1:100 in 0.9% NaCl, and 5 µL added to wells. The prototroph was added to the Transwell while auxotroph added to the main well. Cultures were grown and 75 µL was taken from wells at timepoint for OD600 measurements. For isolate-substrate combinations where we observed slow growth rates, 24, 48 and 72-hour timepoints were used, otherwise timepoints at 0, 24 and 36 hours were used. OD600 values were blanked using no-isolate media controls. For log2 fold-change analysis, values of 0 were transformed to 1x10-5, but kept raw for other statistical analyses. After co-culture, isolates were then plated onto LB agar, confirmed uniform, distinct colony morphology, then re-grown under the same nutrient conditions in single culture.

### Supplemental Results

##### Total number of substrates predicted to support growth varies by strain

Individual *Kp*SC differed in terms of the total number of substrates predicted to support growth, with a general trend towards fewer substrates supported in anaerobic compared to aerobic conditions (262-379 each for aerobic vs 4-320 anaerobic). There was greater variability among usage of carbon sources (104-210 aerobic, 2-173 anaerobic) compared to those used as sources of nitrogen (87-109 aerobic, 2-91 anaerobic), phosphorous (36-52 aerobic, 0-52 anaerobic), and sulfur (7-14 aerobic, 0-10 anaerobic) (**Fig S1A**). This likely reflects a genuine biological trend underpinned by greater variability of carbon metabolic processes as well as biases in our underlying knowledge about metabolism (i.e. carbon metabolism is the most well understood). There were 17 outlier strains which produced predicted growth in only four substrates under anerobic conditions. These were made up of 13 *K. pneumoniae*, 2 *K. quasipneumoniae subsp. similipneumoniae* and 2 *K. variicola subsp. tropica* of various sub-lineages, but were all from the same study (6). This is likely a DNA sequencing artefact as the Nextera XT library preparation was used in this study, which can cause coverage bias issues (7) and likely contributed to Bactabolize not being able to find some key metabolic genes.

#### Substrate usage predictions support species differentiation

As expected, our growth predictions suggested that *Kp*SC taxa are differentiated by core substrate usage patterns (**Fig. S2, Table S5**). Four taxa were represented by sufficient genomes for comparisons (n>200 each); *K. pneumoniae* (hereafter *Kp,* n=6,652), *Klebsiella quasipneumoniae* subsp. *quasipneumoniae* (*Kqq*, n=201), *K. quasipneumoniae* subsp. *similipneumoniae* (*Kqs*, n=285) and *Klebsiella variicola* subsp. *variicola* (*Kvv,* n=672). Among these taxa, a total of 52 distinct growth conditions (corresponding to 18 distinct substrates) were core to at least one but variable and/or absent (not predicted to support growth of any isolates) from at least one other taxon. This included 31 growth conditions that were uniquely core to a single taxon (range 0-16).

All taxa appeared to dedicate similar proportions of their genomes to metabolic orthologs (51.3-54%, **Table S1**), but there were statistically significant differences in the proportions assigned to key macromolecular functions as determined by analysis of Clusters of Orthologous Gene (COG) categories (**Fig. S4**). Notably, *Kp* appeared to have a larger portion of its genes involved with carbohydrate metabolism (median 13.2% vs 12.6-12.9%, $p < 0.0001$ for all comparisons) compared to the other well sampled taxa, and a smaller portion in amino acid transport and metabolism (median 10.04% vs 10.45-10.56%, $p < 0.0001$ for all comparisons). *Kvv* has a smaller portion involved in nucleotide transport and metabolism (median 2.38% vs 2.5-2.53%, $p < 0.0001$), and a larger portion in amino acid metabolism (median 10.56% vs 10.04-10.49%, $p < 0.0001$ for all comparisons).

When compared to the biochemical tests used for formal species definitions, the growth predictions were generally consistent. However, there were several cases that we defined as minor discrepancies where biochemical testing indicated complete absence or conservation of a capability within the KpSC or a specific taxon, and our predictions indicated <5% or >95% conservation, respectively (**Table S6**). These differences likely reflect rare strain variations that have been captured by our genome collection but were not present among the biochemical test data. Seventeen larger-scale discrepancies were also identified; two may reflect inaccuracies in the model predictions for L-carnitine usage, which we previously estimated at ≤30% accuracy (5). Three discrepancies were associated with D-lactic acid methyl ester for which we currently have no prediction accuracy estimate, but for which the biochemically-derived and predicted conservation levels were consistent for taxa represented by ≥200 genomes each. The final 12 discrepancies were associated with substrates for which we have previously confirmed high predictive accuracy from metabolic models (≥94.6%, **Table S6**). It is most likely that these discrepancies were driven by under-sampling in the biochemical testing set, resulting in inaccurate conservation estimates and indicating that the formal species definitions should be revisited.

#### Species and sub-lineages show unique metabolic ortholog fingerprints

Predicted growth phenotype variability is driven by variation in gene and reaction content in the metabolic models, but these represent only the subset of true metabolic genes for which the relevant reaction stoichiometries and supporting literature evidence is available for inclusion in the reference model (5). Therefore, we also explored the distribution of the broader set of metabolic orthologs which showed that each taxon was associated with a distinct core metabolic profile (**Fig. S3, Table S4**). Individual taxa harboured between 145 and 207 core metabolic orthologs beyond the 1,375 that were core to the species complex as a whole. Among the four well sampled taxa (n>200) 63 metabolic orthologs were uniquely core to a single taxon (and absent or variably present in other taxa), whereas 40 were core to at least two taxa and absent or variably present in the others (**Fig. S3, Table S4**).

Within *Kp*, the 48 common sub-lineages were associated with 1,489-1,592 core metabolic orthologs each, plus 75-416 variable orthologs (**Table S4**). Overall, 51 metabolic orthologs were majority sub-lineage specific core (core to ≥66% of sub-lineages); 56 orthologs were common sub-lineages specific core (≥25% to <66% of sub-lineages) and 127 were rare sub-lineage specific core. 888 metabolic orthologs were not core to any sub-lineage but variably present in between 0 and 48 well-represented sub-lineages.

#### Co-occurrence analysis of metabolic traits

Co-occurrence analysis was performed on growth phenotype predictions to identify genetically linked and/or co-selected traits. Isolates were subsampled to a maximum of 10 genomes per sub-lineage to control for population sampling biases (n=2,427 genomes, **Table S1**). Twelve pairs of growth phenotypes (substrate usage) were detected as co-occurring in this dataset including five instances of a single substrate utilised as alternative element sources (e.g. carbon and nitrogen) plus four additional mechanistically linked substrate pairs: Methanesulfonate (as a sulfur source) and methanol usage were linked by their converging degradation into formaldehyde, though methanesulfonate additionally produces sulfite as a sulfur source. Galactitol and D-tagatose usage were linked by their common degradation into D-tagatofuranose 1,6-bisphosphate by two distinct enzymes (tagatose-bisphosphate aldolase and phosphofructokinase). Fructoselysine and xylitol usage were linked by their converging degradation into D-ribulose 5-phosphate, a key substrate for the pentose phosphate pathway and an essential substrate for biomass production. Acetate and butyrate are closely linked by their direct degradation into butanoyl-CoA, an important co-enzyme involved in fatty acid metabolism. In contrast, for the twelfth pair formamide and Fe(III)dicitrate, we could not identify a common mechanistic pathway or enzyme, indicating co-occurrence may be due to co-selection or physical genetic linkage.

#### Experimental validation of growth predictions

*In vitro* growth assays were performed on seven substrates as sole sources of carbon plus allantoin as a sole source of nitrogen and methanesulfonate as a sole source of sulfur in m9 minimal media, aerobic atmosphere. These were selected to represent common and rare sub-lineage specific core growth capabilities that had not been validated previously (5) as well as two positive controls that are known to support growth of all *K. pneumoniae* (glucose and glycerol). Two experiments were performed, i) measuring $OD_{600}$ at 24- and 48-hour endpoints; ii) a substrate coaxing experiment (see **Methods**). This second experiment was performed for substrates where one or more metabolic model predicted growth but growth was not observed in the initial 48-hour culture assays. These types of false positive model predictions are most likely due to regulatory mechanisms that are not considered by the models and may indicate that the isolates were not expressing relevant genes. To address this, isolates were coaxed onto the testing substrate via subculturing into progressively lower amounts of glycerol (core carbon source; 10 mM on subculture 1, 5 mM on subculture 2, 0 mM on subculture 3) plus the substrate of interest.

A total of 13 distinct *K. pneumoniae* isolates, representing nine distinct sub-lineages, were tested in triplicate, grown aerobically on each of the nine substrates (**Table S2**)**.** As expected, all isolates were able to grow in M9 plus glucose and M9 plus glycerol. Similarly, all isolates that were predicted to grow in M9 plus L-hydroxyproline were able to do so, as were two additional isolates (AJ229, INF269) that were not predicted to be able to utilise this substrate. Unexpectedly, acetoacetate, allantoin and methanesulfonate supported growth of 12/13, 13/13 and 12/13, isolates respectively, including 10, 11 and nine isolates that were not predicted to grow. These results indicate that the metabolic models are missing genes and reactions that support the metabolism of these substrates, but database and literature searches did not resolve these issues.

Growth on allantoin as a nitrogen source was particularly interesting as it has been previously shown to be specific and in-fact essential for virulence in *K. pneumoniae* SL23 (8). Further investigation showed that the necessary reactions to support allantoin metabolism were present in all models, but the transport reaction, required for transport of the substrate into the cytosol, was missing in most. The exceptions were models carrying the *all* accessory operon including those from SL23 (**Fig. 3**). However, our *in vitro* coaxing experiments indicate that all *K. pneumoniae* may be able to utilise this substrate to some degree given the right conditions. As our experiments grew to three days at 60 mM allantoin (considerably higher concentration than tested previously (8)), we were able to detect this growth. We predict that SL23 may utilise allantoin more efficiently, due to the presence of the dedicated transport machinery, but further investigations are needed (beyond the scope of this work).

In contrast to those discussed above, we were not able to demonstrate growth of any isolates on butyrate or methanol as sole sources of carbon. We suspected that we did not capture the correct growth conditions for these substrates, as the genes required for their metabolism were clearly present within some of the test isolate genomes. Notably, previously generated growth profiles of 37 isolates in aerobic (9) and anaerobic conditions (5), indicated that *Kp*SC prefer anerobic usage of butyrate; 17/37 isolates were able to grow on butyrate resulting in a model predictive accuracy of 94.59%.

Finally, the *in vitro* data indicated several inaccuracies for prediction of growth on xylitol, with three false positive and four false negative model predictions for the 13 tested isolates.

#### *In silico* co-culture experiments

In order to better understand the interactions between the auxotroph and prototroph isolate pairs tested *in vitro*, we simulated co-cultures *in silico* using MICOM (10). This approach aims to optimise growth of the community (the isolate pairs in this case) and will therefore favour mutualistic interactions. Since we had already demonstrated such interactions *in vitro*, we were seeking to identify the set of putative cross-feeding metabolites, i.e. those with the highest flux donated from the prototroph to auxotroph.

For Galactitol, L-hydroxyproline, L-sorbose and L-tartrate as sole carbon sources, the metabolites with the lowest negative flux from the prototroph (produced by) and the highest positive flux to the auxotroph (gained by) were identified. These included acetaldehyde, (R)-glycerate, acetate, ethanol, 4-aminobenzoyl-glutamate, L-glutamate and pyruvate. Each of these metabolites supported growth as sole carbon sources for the auxotroph. This mechanism likely explains the growth of auxotrophs in co-culture, whereby, the donor prototroph catabolises the substrate, and exports metabolites that the auxotroph is capable of utilising as a carbon source.

### Supplementary Tables

**Table S1 (Excel file):** List of all 7,835 genomes used for this study along with accessions, metadata, per genome summary of simulated growth phenotypes and proportional Clusters of Orthologous Genes (COG) data.

**Table S2 (Excel file):** Outcomes of *in vitro* growth experiments for metabolic model validation.

**Table S3 (Excel file):** Raw data and statistical analysis of co-culture experiments.

**Table S4 (Excel file):** Metabolic gene (metabolic KEGG Ortholog) presence/absence table for entire 7,835 genome dataset.

**Table S5 (Excel file):** Binarised growth predictions from metabolic modelling for entire 7,835 genome dataset.

**Table S6:** Biochemical tests described in formal species definitions and consistency with model predictions.

| Biochemical test | Taxa (phylogroup) | | | | | | | Growth prediction accuracy (%) (5) |
|---|---|---|---|---|---|---|---|---|
| | *Kp* (Kp1) | *Kqq* (Kp2) | *Kvv* (Kp3) | *Kqs* (Kp4) | *Kvt* (Kp5) | *Kqv* (Kp6) | *Ka* (Kp7) | |
| **Substrates for which growth was thought to be conserved among KpSC (11)** | | | | | | | | |
| D-Glucose | (100) | (100) | (100) | (100) | (100) | (100) | (100) | 100 |
| L-Arabinose | (100) | (100) | (100) | (100) | (100) | (100) | (100) | 100 |
| D-Mannitol | (100) | (100) | (100) | (100) | (100) | (100) | (100) | 92 |
| D-Mannose | (100) | (100) | (100) | (98) | (100) | (100) | (100) | 100 |
| Melibiose | (100) | (100) | (100) | (98) | **(60)** | (100) | (100) | 100 |
| Raffinose | (100) | (100) | (100) | (100) | (100) | (100) | (100) | 100 |
| L-Rhamnose | (100) | (100) | (100) | (100) | (100) | (100) | (100) | 97 |
| Salicin | (100) | (100) | (100) | (100) | (100) | (100) | (100) | 100 |
| D-Sorbitol | (100) | (100) | (100) | (98) | (100) | (100) | (100) | 100 |
| Sucrose | (99) | (100) | (100) | (99) | (100) | (100) | (100) | 100 |
| Trehalose | (100) | (100) | (100) | (100) | (100) | (100) | (100) | 100 |
| D-Xylose | (100) | (100) | (100) | (100) | (100) | (100) | (100) | 78 |
| D-Arabitol | (99) | (100) | (99) | (100) | (100) | (100) | **(0)** | 97 |
| Cellobiose | (100) | (100) | (100) | (100) | (100) | (100) | (100) | 100 |
| Lactose | (100) | (100) | (100) | (100) | (100) | (100) | (100) | 100 |

| | | | | | | | | |
|---|---|---|---|---|---|---|---|---|
| Maltose | (97) | (100) | (100) | (98) | (100) | (100) | **(0)** | 95 |
| Glycerol | (100) | (100) | (100) | (100) | (100) | (100) | (100) | 97 |
| Urea (nitrogen) | (100) | (100) | (100) | (100) | (100) | (100) | (100) | Not tested |
| L-Tartrate | **(67)** | **(74)** | (100) | **(30)** | (100) | (100) | (100) | 95 |
| myo-Inositol | (100) | (100) | (100) | (98) | (100) | (100) | (100) | 100 |
| **Substrates that differentiate KpSC taxa (12)** | | | | | | | | |
| Dulcitol | ~ (48) | - (1) | ~ (58) | ~ (34) | ~ (80) | + (100) | + (100) | 100.0 |
| Adonitol | + (99) | ~ (63) | - (1) | + (99) | - (0) | - (0) | - (0) | 100.0 |
| Tricarballylic acid | - (0) | + (100) | ~ (100) | + (97) | + (60) | ~ (93) | + (100) | 94.6 |
| N-acetyl-neuraminic acid | - (0) | - (0) | - (0) | + (97) | - (0) | - (0) | - (0) | 100.0 |
| D-arabitol | + (99) | + (100) | + (99) | + (100) | + (100) | + (100) | - (0) | 100.0 |
| L-sorbose | ~ (53) | - (33) | + (100) | - (0) | + (100) | + (100) | + (100) | 97.3 |
| D-tagatose | ~ (48) | - (1) | ~ (58) | ~ (34) | ~ (80) | + (100) | + (100) | 100.0 |
| 5-keto-D-gluconic acid | - (0) | ~ (41) | + (100) | - (3) | + (100) | + (100) | + (100) | 100.0 |
| D-lactic acid methyl ester | + (100) | + (100) | + (100) | + (99) | - (100) | - (100) | - (100) | Not tested |
| 4-hydroxyl-L-proline | - (11) | + (75) | + (100) | + (98) | + (100) | + (100) | + (29) | 100.0 |
| L-carnitine | ~ (0) | - (0) | + (0) | + (0) | + (0) | - (0) | + (0) | 29.7 |
| **Sample sizes** | | | | | | | | |
| N isolates (biochemical tests) (12) | 10 | 5 | 5 | 4 | 6 | 5 | 1 | N/A |
| N genomes (growth predictions) (This study) | 6652 | 201 | 672 | 285 | 5 | 13 | 7 | N/A |

Substrate usage conservation as determined by biochemical testing reported in (11, 12) and predicted in this study (parentheses) is shown for each KpSC taxon: Kp = *K. pneumoniae,* Kqq = K. *quasipneumoniae* subsp. *quasipneumoniae*, Kvv = *K. variicola* subsp. *variicola,* Kqs = K. *quasipneumoniae* subsp. *similipneumoniae*, Kvt = *K. variicola* subsp. *tropica*, Kqv = *K. quasivariicola,* Ka = *K. africana* (labels in parentheses are phylogroup designations reported in (11)). '+' indicates 100% conservation in biochemical testing, '-' indicates 0% conservation in biochemical testing, '~' indicates >0% and <100% conservation in biochemical testing (absolute values not reported). Values in parentheses indicate the percentage of isolates predicted to be able to utilise the substrate in this work. Formatting indicates the level of concordance between the conservation values derived from biochemical tests and those derived from growth predictions; none = conservation levels are consistent; underline = minor discrepancies i.e. predicted conservation level <5% different from biochemical conservation level (applied only where biochemical testing indicated 0% or 100% conservation); bold and grey shading = conservation levels are not consistent. Substrate level growth prediction accuracies are also indicated, as reported in (5).

### Supplementary Figures

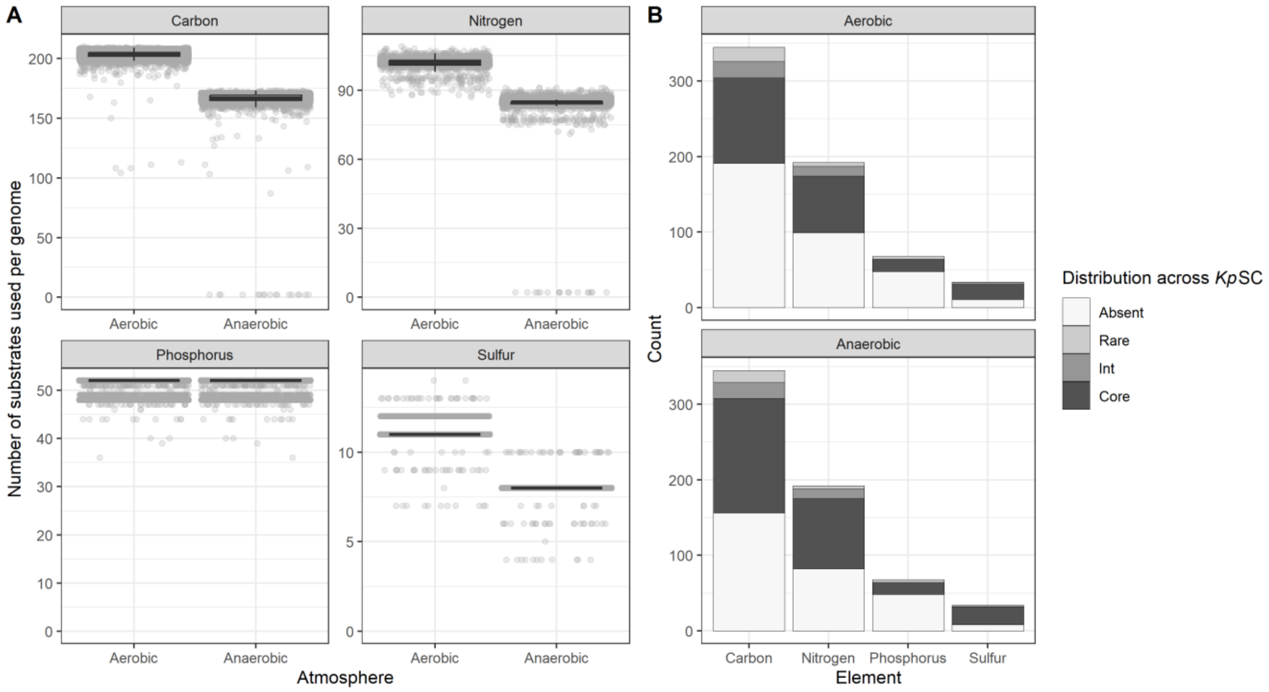


**Fig. S1: Substrate usage frequencies across *Kp*SC dataset**

**A:** Number of substrates predicted to support growth for each isolate, stratified by substrate type and atmosphere. **B:** Population frequency of predicted substrate usage by substrate type. Core refers to substrate usage conserved in ≥95% of isolates. ‘Int’ refers to intermediately present (>15% to <95%). Rare refers to >0 to <15%, while ‘Absent’ indicates no predicted growth. All binarised growth predictions can be found in **Table S3**.

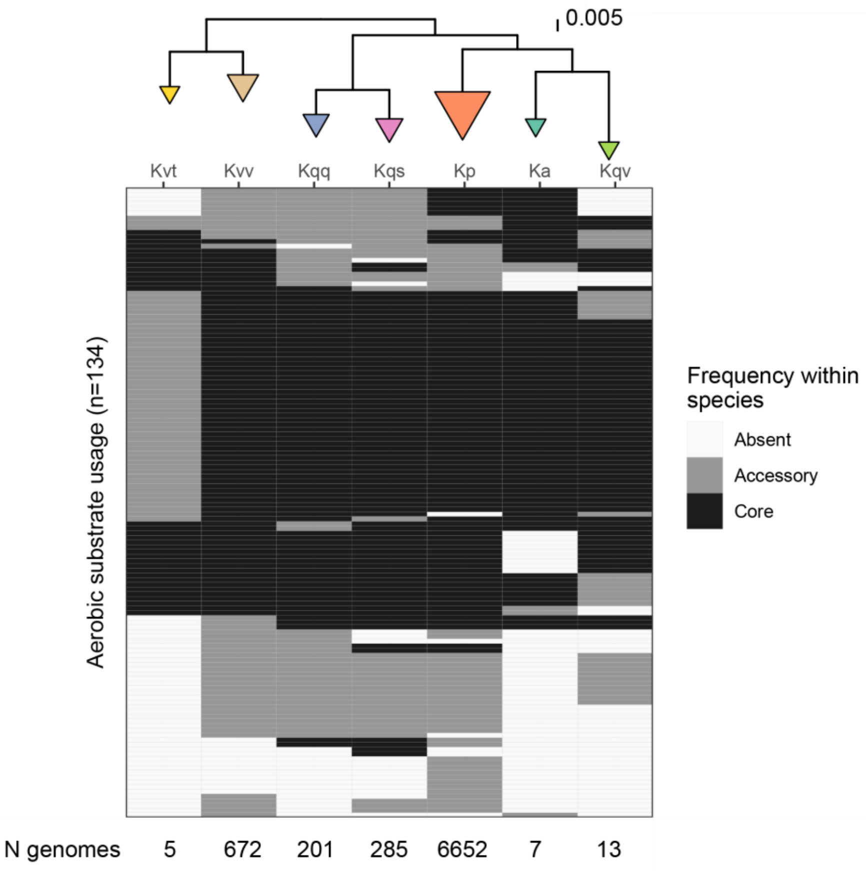


**Fig. S2: Taxon-specific substrate usage**

Heatmap showing taxon-specific substrate usage as predicted using metabolic models. Only substrates for which usage frequencies differed in aerobic conditions are shown. Columns are arranged by species-phylogeny and the number of genomes per taxon is shown along the bottom of the x-axis and arrowhead size. Taxa names shortened for brevity: Ka = *K. africana*. Kp = *K. pneumoniae*. Kqq = K. *quasipneumoniae* subsp. *quasipneumoniae*. Kqs = K. *quasipneumoniae* subsp. *similipneumoniae*. Kqv = *K. quasivariicola*. Kvt = *K. variicola* subsp. *tropica*. Kvv = *K. variicola* subsp. *variicola*.

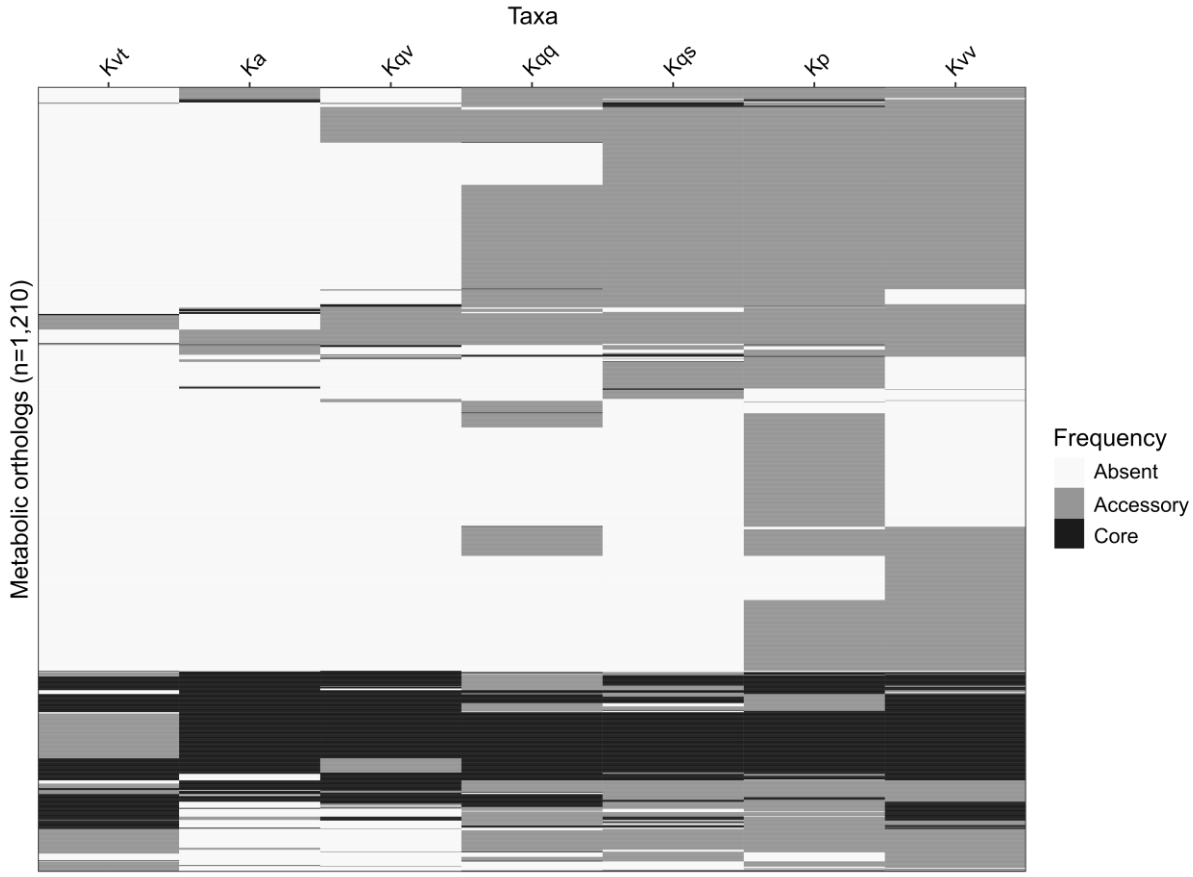


**Fig. S3: Taxa are associated with unique metabolic ortholog profiles**

Heatmap showing taxon-specific metabolic orthologs. Only orthologs for which frequencies differed between taxa are shown. Columns are arranged by species-phylogeny as in **Fig. S2** and names shortened for brevity: Ka = *K. africana*. Kp = *K. pneumoniae*. Kqq = K. *quasipneumoniae* subsp. *quasipneumoniae*. Kqs = K. *quasipneumoniae* subsp. *similipneumoniae*. Kqv = *K. quasivariicola*. Kvt = *K. variicola* subsp. *tropica*. Kvv = *K. variicola* subsp. *variicola*. Frequencies are indicated by colours as shown in the legend: Absent = not present in any genomes; Accessory = present in >0% and <95% genomes; Core = present in ≥95% genomes. Raw data can be found in Table S2

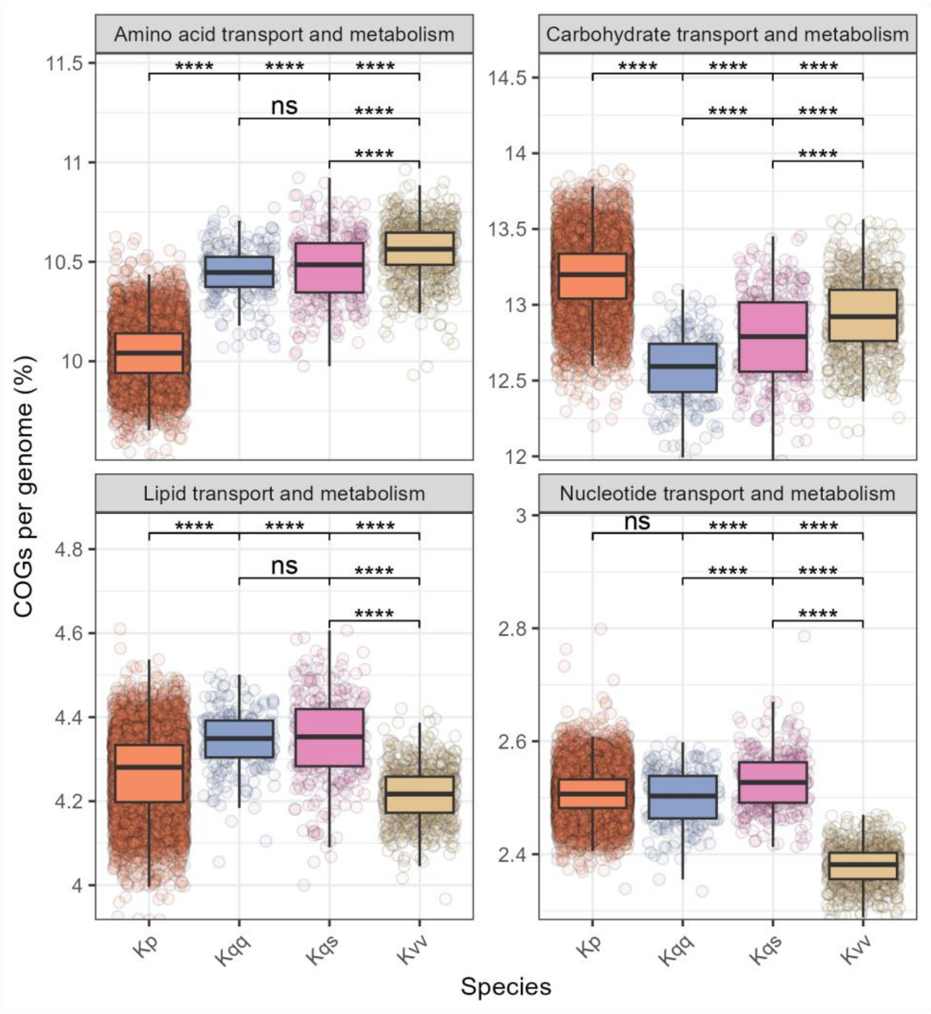


**Fig. S4: Taxon-specific specialisation of macromolecular functions**

Box and whisker array showing the proportion of each genome assigned to Clusters of Orthologous Gene categories (COG) by taxon coloured as per **Fig. S2**. Dots show individual genomes. Significance is indicated such that the first row shows all comparisons to *K. pneumoniae* (Kp), the second row shows comparisons to *K. quasipneumoniae* subsp. *quasipneumoniae* (Kqq) and third row shows comparisons between *K. quasipneumoniae* subsp. *similipneumoniae* (Kqs) and *K. variicola* subsp. variicola (Kvv). Significance calculated using a non-parametric Kruskal–Wallis test with Holm correction, followed by Dunn's *post-hoc* test (****: $p < 0.0001$, ns: not significant). Only species with n>200 genomes were analysed.

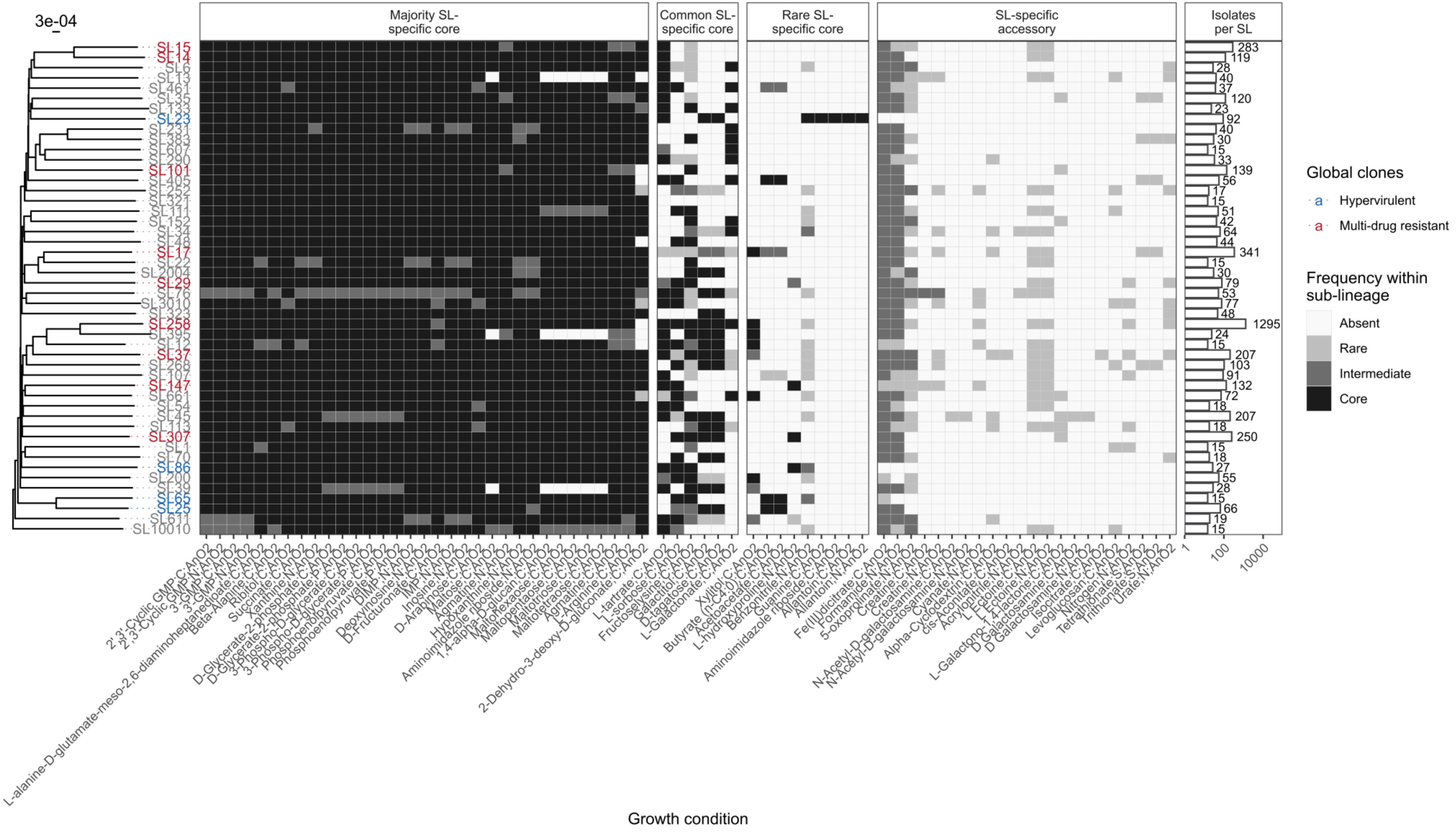

3e-04
Majority SL-specific core
Common SL-specific core
Rare SL-specific core
SL-specific accessory
Isolates per SL
Global clones
Hypervirulent
Multi-drug resistant
Frequency within sub-lineage
Absent
Rare
Intermediate
Core
Growth condition

**Fig. S5: Distinct anerobic metabolic fingerprints of sub-lineages within species**.

Heatmap showing frequency of variable anaerobic substrate usage across 48 global *K. pneumoniae* sub-lineages (SL). Rows are ordered by phylogeny. Sub-lineage labels are coloured to indicate the globally-distributed clones described in (13): Blue shows hypervirulent while red shows multidrug resistant. Substrate shown along X-axis. The element source of each substrate is semi-colon separated and abbreviated for brevity: C = Carbon, N = Nitrogen and S = Sulfur. AnO2 indicates anaerobic conditions. Frequency of substrate usage indicated by shading as shown in legend. Number of isolates per each sub-lineage shown in bars.
