## Supplementary figures and images for "A metabolic atlas of the *Klebsiella pneumoniae* species complex reveals lineage-specific metabolism that supports co-existence of diverse lineages"

### Fig. S1

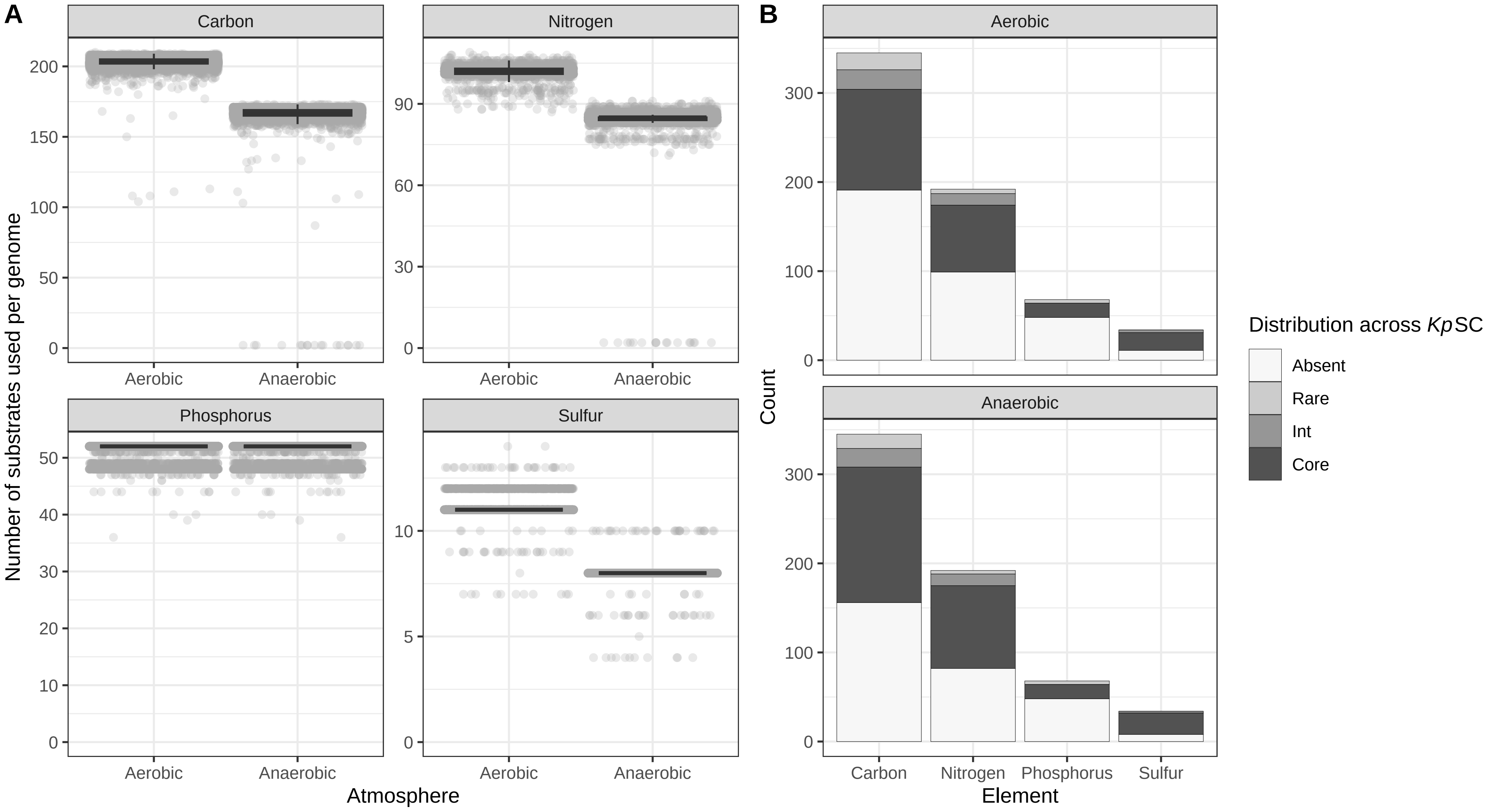

### Fig. S2

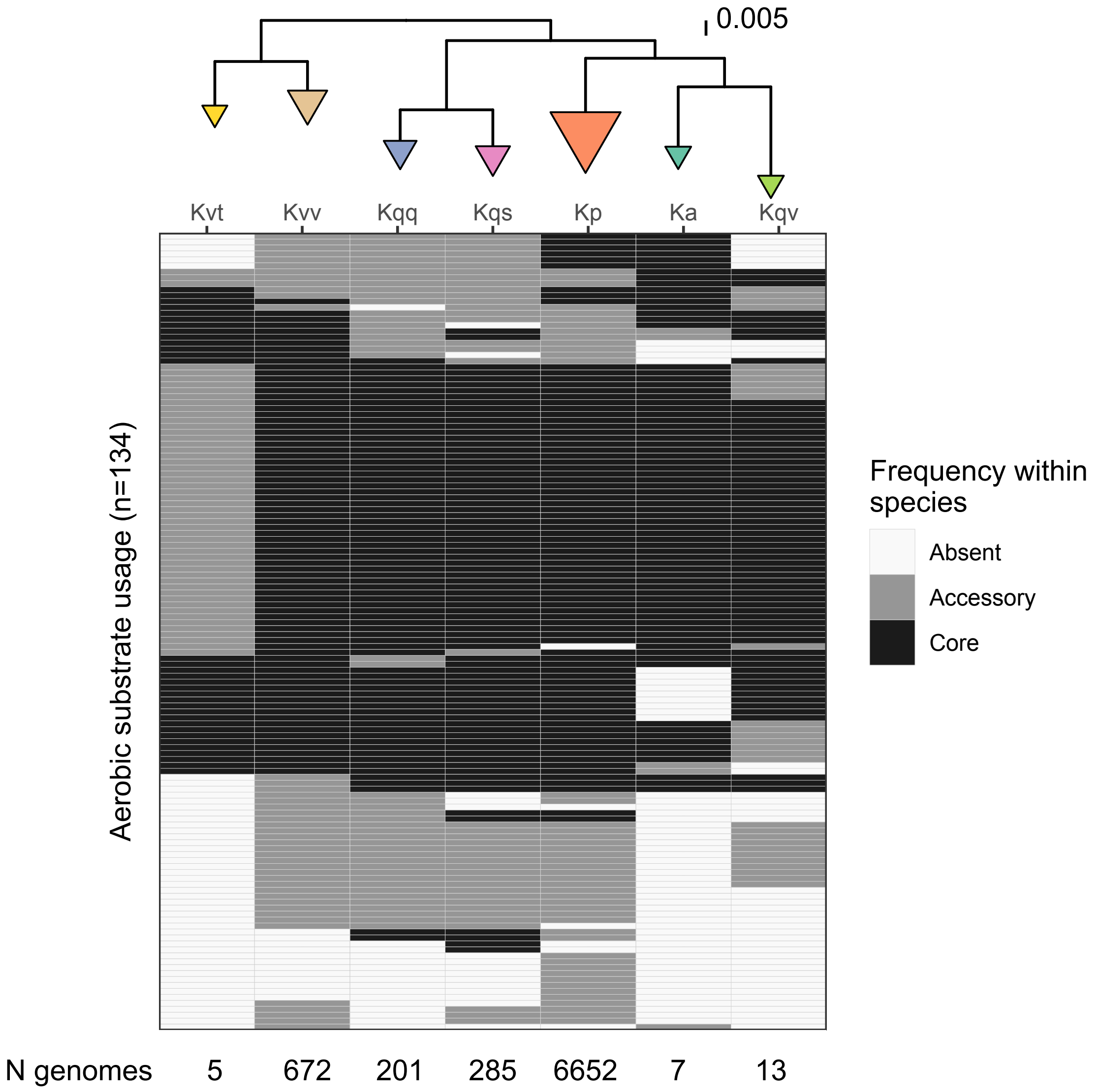

### Fig. S3

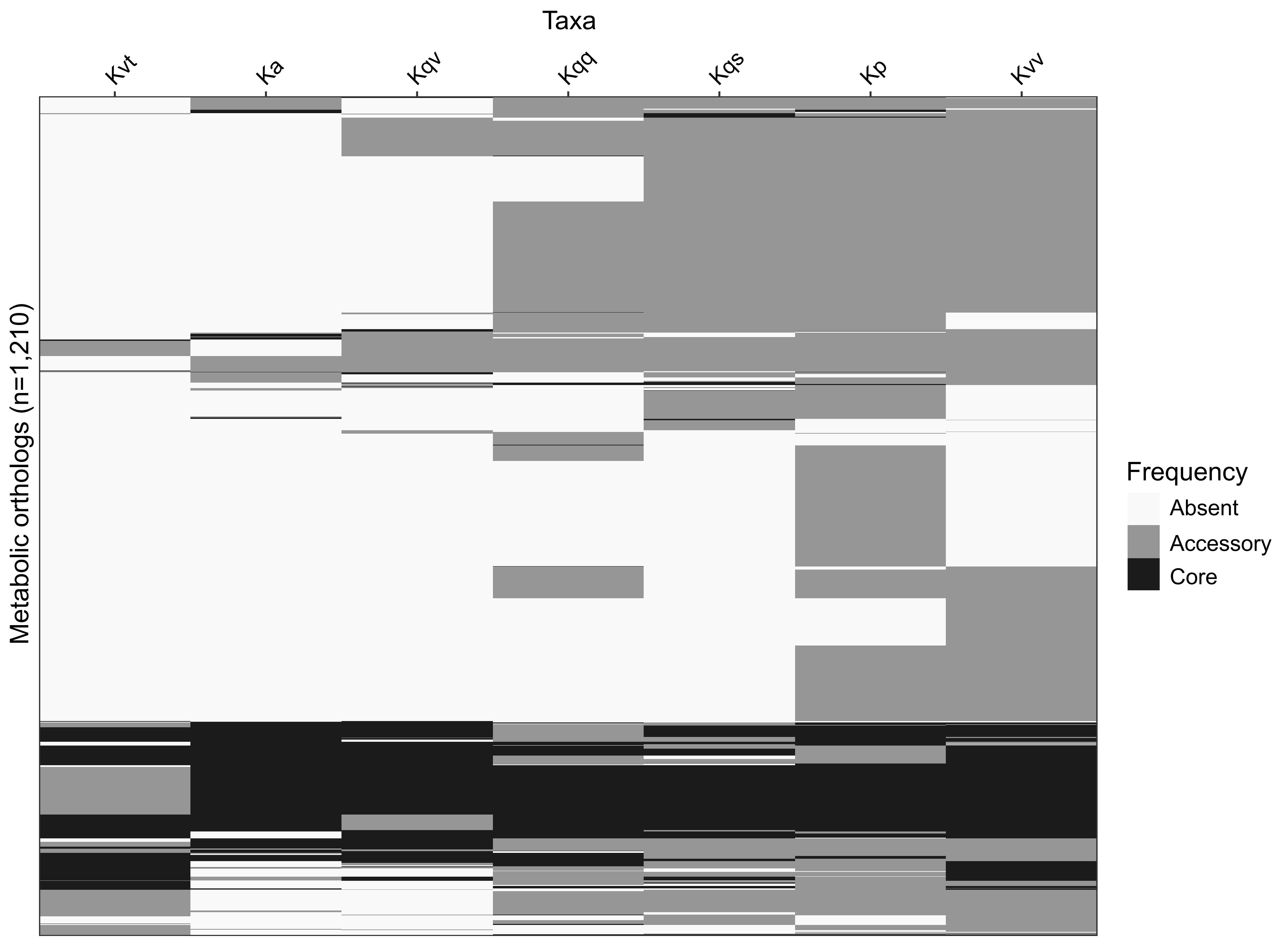

### Fig. S4

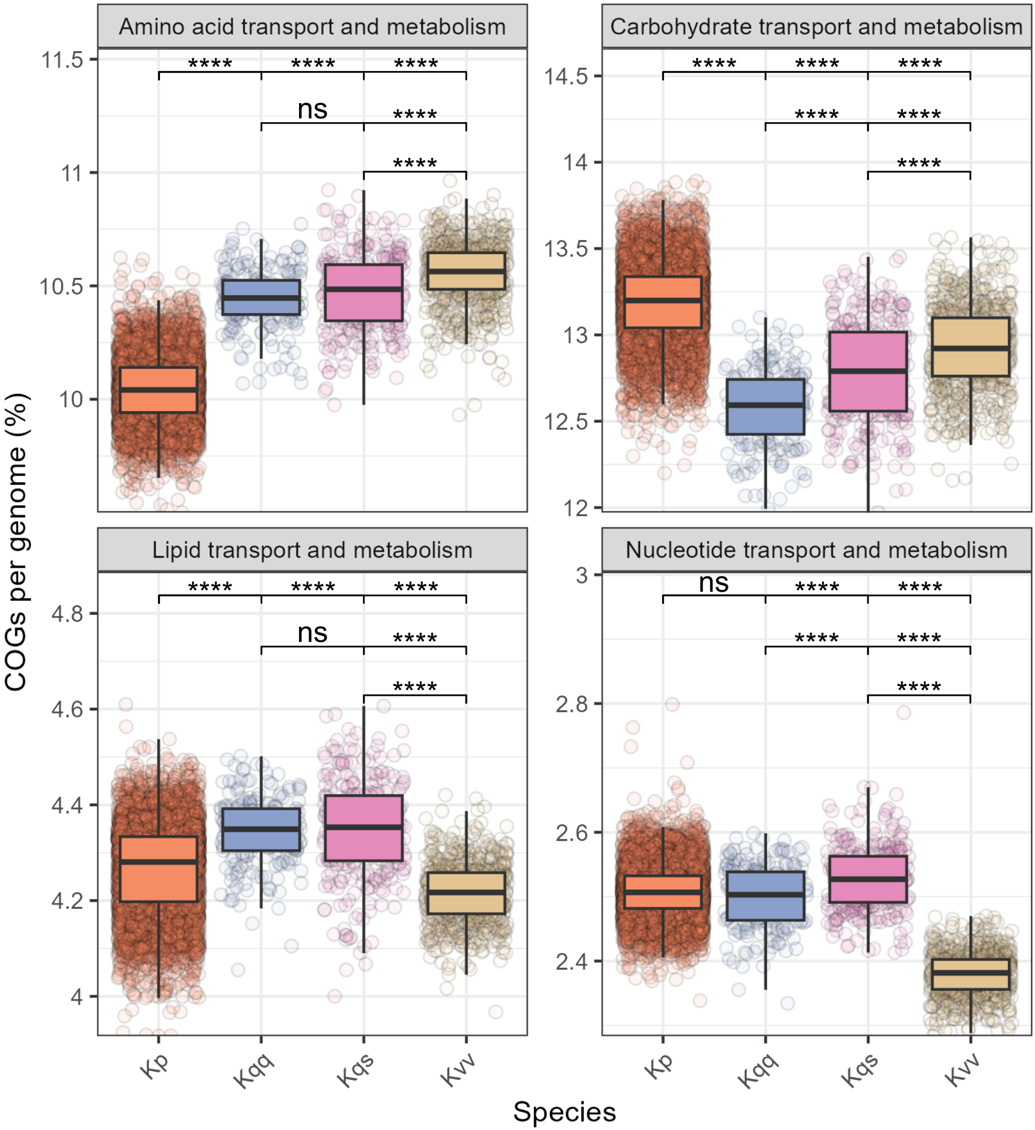

### Fig. S5

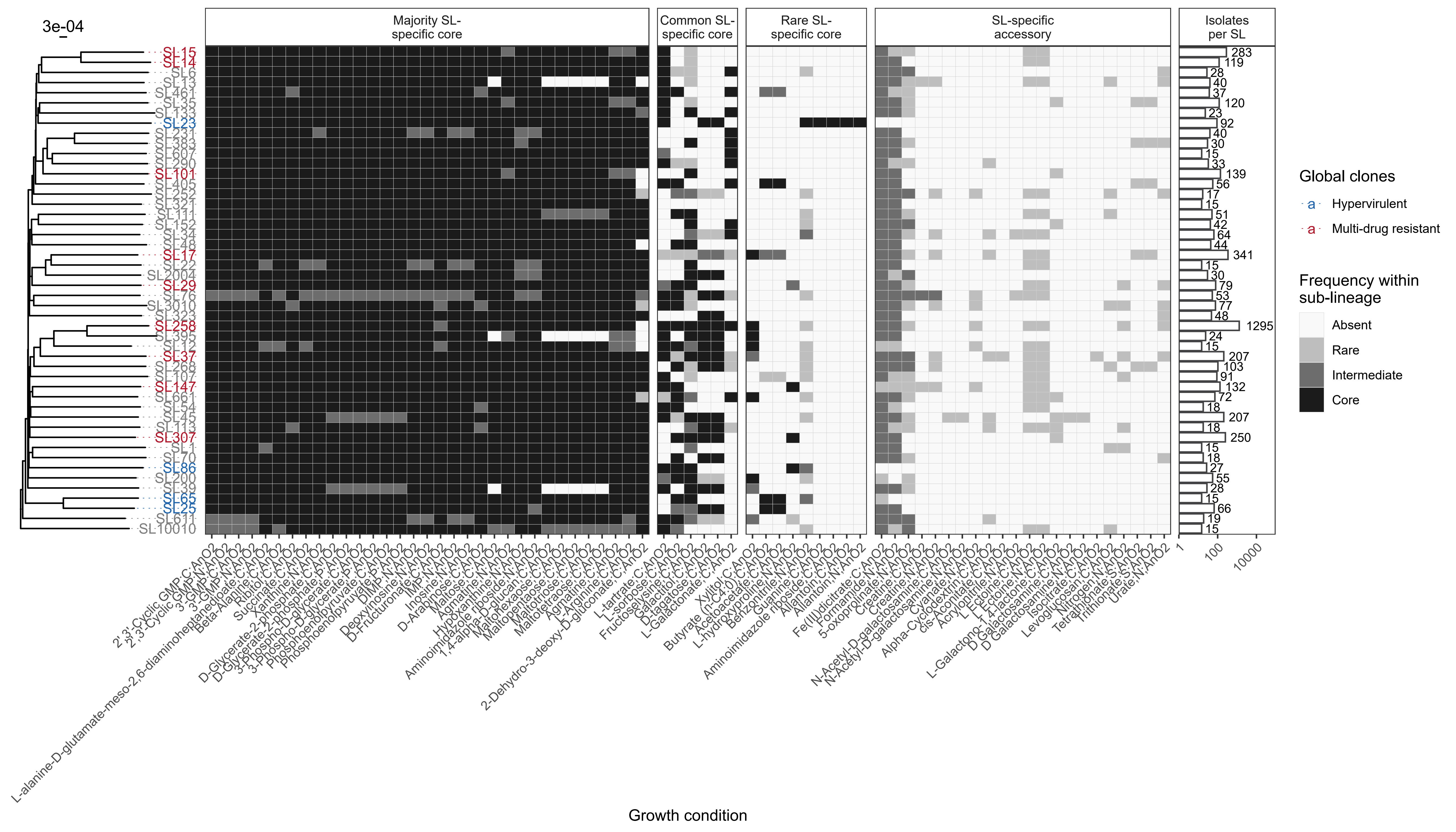
